# Impaired envelope integrity in the absence of SanA is linked to increased lipid II availability and an imbalance of septal peptidoglycan synthesis

**DOI:** 10.1101/2025.06.10.658892

**Authors:** Joseph F. Carr, Carolina Basurto De Santiago, Shivani A. Bhut, Daniel J. Warzecha, Sarah A. Vastani, Robert Wei, Carmen M. Herrera, M. Stephen Trent, Beiyan Nan, Angela M. Mitchell

## Abstract

In Gram-negative bacteria, the outer membrane (OM) acts in conjunction with the peptidoglycan (PG) cell wall as a barrier against physical, osmotic, and chemical environmental stressors including antibiotics. SanA, an inner membrane protein in *Escherichia coli* K-12, is required for vancomycin resistance at high temperatures (>42 °C) and impacts sodium dodecyl sulfate (SDS) resistance during stationary phase reached from carbon limitation. However, its function remains unknown. Here, we show that Δ*sanA* has a synthetic genetic interaction with Δ*wecA*, a mutation that increases the availability of the isoprenoid carrier for PG synthesis. Specifically, the Δ*sanA* Δ*wecA* strain demonstrated heightened SDS-EDTA sensitivity, activation of the Rcs stress response, and increased cell length. Further investigation tied the SDS-EDTA sensitivity to increased lipid II available for PG synthesis. Spontaneous suppressor mutants of this phenotype harbored point mutations in *prc*, which encodes tail specific protease, or *ftsI*, which encodes the cell division DD-transpeptidase, a target of Prc. We focused on the *ftsI* mutations and demonstrated that the *ftsI* mutations had increased cell length but nevertheless enhanced PG incorporation at the septum compared to the Δ*sanA* mutant, returning PG incorporation to wild-type levels. Moreover, other mutations affecting septal PG synthesis, but not divisome assembly, also suppressed the SDS-EDTA sensitivity. These findings suggest that, in the absence of SanA, increased lipid II availability perturbs the balance between septal PG synthesis, lateral PG elongation, and other envelope biogenesis pathways, which leads to increased OM permeability.

**IMPORTANCE:** The Gram-negative cell envelope is a barrier that protects the cell from environmental stress. Therefore, the synthesis of each layer of this envelope needs to be closely coordinated throughout growth and division. Here, we investigated SanA, a protein in *Escherichia coli* K-12 that affects envelope permeability under cellular stress, including nutrient limitation and high temperature. We found that SanA plays a key role in maintaining the permeability barrier when precursor levels for peptidoglycan (PG) synthesis are elevated, linking envelope integrity to balanced septal PG production during cell division. Our results suggest that SanA modulates substrate availability to preserve envelope function, and that in its absence, imbalanced substrate flux to septal PG synthesis disrupts septum formation and compromises barrier integrity.

## INTRODUCTION

Gram-negative bacteria have a multi-layered envelope consisting of the inner or cytoplasmic membrane (IM), the aqueous periplasm containing the peptidoglycan (PG) cell wall, and the outer membrane (OM) (1). While the IM is a phospholipid bilayer, the lipid portion of the OM is composed of phospholipids in the inner leaflet and primarily lipopolysaccharide (LPS) in the outer leaflet. The cell envelope is tightly interconnected. For instance, the OM and PG are physically connected by an OM lipoprotein (Lpp) and/or outer membrane proteins (OMPs) that bind to PG (e.g., OmpA) (2, 3). Properly composed, the cell envelope prevents entry of toxic molecules, resists osmotic and physical stress, and facilitates cell growth, all while allowing nutrients to enter the cell and metabolic waste to be expelled (1, 4–8). The envelope prevents entry of molecules that are either large or hydrophobic, making Gram-negative bacteria intrinsically resistant to many clinically used antibiotics and greatly complicating discovery of new antimicrobials (4). Yet, the pathways responsible for envelope biogenesis and the regulatory interactions between the biogenesis pathways are promising targets for development of new antimicrobials (9–17).

While synthesis of each envelope component depends on a separate pathway (1, 18–21), complex regulatory interactions between these pathways are essential to coordinate growth of the layers (22). For instance, in *Escherichia coli*, halting fatty acid synthesis, required for phospholipid and LPS synthesis, also causes PG synthesis to cease (23). Conversely, excess phospholipid synthesis can trigger PG expansion. In this case, phospholipids in the outer leaflet of the OM act as a signal to activate MepS, an endopeptidase that breaks PG crosslinks to allow new PG insertion (24). Interestingly, cell growth can be separated from new PG synthesis and endopeptidase activity, which can continue in the absence of PG synthesis until the point of cell lysis (25). YhcB (a.k.a. ZapG, LapD) has been suggested to interact with both the elongasome (the machinery responsible for PG synthesis during cell growth) and with the complex responsible for regulating LPS synthesis and to play a role in balancing the production of LPS, PG, and phospholipids (26–28). However, recent results have demonstrated that YhcB interacts directly with AccA, a member of the complex that initiates fatty acid synthesis, and the phenotypes of a *yhcB* mutant are directly tied to an increase in phospholipid production due to excess fatty acid synthesis (29, 30). This suggests the effects on PG and LPS production are a secondary consequence of the complex regulatory interactions between cell envelope pathways. The OM’s physical strength also plays a role in orienting PG synthesis, and LPS mutations strengthening the OM can restore rod shape in mutants deficient in activation of the elongasome (31). In *Pseudomonas aeruginosa*, MurA, which catalyzes the first committed step in PG biosynthesis, directly interacts with LpxC, which catalyzes the first committed step in LPS biosynthesis, causing stimulation of LpxC activity (32).

Interactions in biosynthesis can also occur through shared synthesis substrates. For instance, most extracytoplasmic glycans are synthesized on an isoprenoid carrier, undecaprenyl-phosphate (Und-P), and mutations that affect biosynthesis of one glycan can disrupt synthesis of others due to sequestration of Und-P. Enterobacterial common antigen (ECA) is an extracytoplasmic glycan found in the Enterobacterales envelope and consists of a repeating, invariant trisaccharide unit that is found surface exposed as the headgroup of a phospholipid or attached to LPS in place of O-antigen, as well as cyclized in the periplasm (33). Mutations in later steps of ECA synthesis cause cell shape defects and envelope stress response activation, presumably through inhibition of PG synthesis (34–39). Loss of WecA, which catalyzes the first step in ECA synthesis, causes an approximately 2-fold increase in free Und-P levels (40). Loss of later steps in ECA biosynthesis cause Und-P levels to decrease compared to Δ*wecA*, with later mutations causing larger decreases.

The need for coordination in synthesis of the cell envelope layers becomes more acute during conditions that perturb synthesis of envelope components such as nutritional stress or cell division. PG synthesis during cell division is carried out by a highly regulated complex known as the divisome (41, 42). The divisome is assembled on the FtsZ ring which consists of FtsZ, FtsA, and ZipA in *E. coli*. FtsK is then recruited, followed by FtsQ, FtsL, and FtsB. Once this complex is complete, FtsW, FtsI, and FtsN then join the complex. FtsA plays a key role in recruiting the downstream components of the complex, while FtsN recruitment acts as a signal to begin septal PG synthesis (41, 42). FtsW and FtsI (PBP3) are responsible for septal PG synthesis with FtsW acting as the transglycosylase linking the glycan backbone of PG and FtsI acting as the DD-transpeptidase to link the peptide stems of the PG.

During septal PG synthesis, the OM invaginates with the PG allowing coordinated separation of the cell envelope (42). This coordination is accomplished through a protein, CpoB, that interacts with LpoB and PBP1B at the septum and with the Tol-Pal complex, which has IM and OM components and contacts PG (43–45). Beyond its importance for OM invagination, Tol-Pal, as well as two OM lipoproteins (NlpD and YraP), are involved in the activation of amidases that split the septal PG during cell division, ensuring the coordination of OM invagination with cell division (46, 47). Interestingly, mutations inactivating NlpD lead to increased sensitivity to the detergent SDS (sodium dodecyl sulfate) in combination with EDTA, which disrupts interactions between LPS molecules, suggesting that the OM permeability barrier is disrupted by loss of cell division amidase function (46). To complete OM invagination, an increase in OM synthesis is required. Folding of OMPs into the OM by the β-barrel assembly machine (BAM) is activated by immature PG allowing insertion of new OMPs into the OM at the cell septum (48), and so allowing balanced OM synthesis during cell division.

We previously investigated changes in OM permeability during stationary phase initiated by carbon limitation (49). We found that resistance to SDS during carbon limitation did not rely on efflux pumps as it did in exponentially growing cells. Instead, a RpoS (the master regulator of stationary phase)-dependent change in OM permeability occurs, which requires the presence of three genes, *sanA*, *dacA* (PBP5), and *yhdP* (49). YhdP is a presumptive inter-membrane phospholipid transporter (50–53), the loss of which causes increased OM permeability in the presence of ECA (54). PBP5 is a carboxypeptidase that cleaves D-alanine from the PG pentapeptide, regulating the amount of substrate available for transpeptidases that crosslink PG (55). Loss of PBP5 can prevent the maturation of PG that results in mid-cell insertion of new OMP proteins (48). Together, these findings suggest that envelope coordination is important for the stationary phase membrane permeability. However, the function of *sanA* has remained unknown.

SanA is a bitopic IM protein with one N-terminal transmembrane helix and a periplasmic DUF218 domain (56, 57). In *E. coli* K-12, *sanA* was first identified as a multicopy suppressor of vancomycin sensitivity for a strain that had a large gene deletion including *sanA* (58). It was also found to be involved in vancomycin resistance at high temperature (i.e., 43 °C) (58). In *Salmonella enterica* serovar Typhimurium, SfiX, a SanA homolog, is necessary for filamentation of a mutant overexpressing *hisFH* (His^C^) and also affects vancomycin resistance at high temperature (59). Recent work demonstrated a decrease in hydrophobicity and charge of the *Salmonella* cell surface and a corresponding change in envelope permeability in a *sfix* (*sanA*) mutant (60). Here, we investigated the envelope permeability phenotypes caused by SanA’s loss and found a genetic interaction with ECA synthesis. Mutations expected to increase availability of Und-P caused the *sanA* mutant to become sensitive to SDS-EDTA and activated the Rcs (Regulator of Capsule Synthesis) envelope stress response. We then selected suppressors of SDS-EDTA sensitivity and isolated partial loss-of-function mutations in *ftsI* (PBP3) and in *prc*, a protease involved in the maturation of FtsI (61, 62). This finding suggests a connection between cell division and OM permeability. Testing of additional cell division mutations demonstrated that mutations in septal PG synthesis but not divisome assembly suppressed the OM permeability defect. Our results tie the envelope permeability phenotype to lipid II availability and support a model in which SanA modulates the availability of lipid II for septal PG synthesis. We propose that, when lipid II levels are high such as during certain cellular stresses, lack of balance between septal PG synthesis and other envelope biogenesis pathways alters OM composition and envelope permeability.

## RESULTS

### Loss of SanA increases membrane permeability under cellular stress conditions

We previously reported that, when treated during stationary phase reached from carbon limitation, a Δ*sanA* strain is sensitive to the detergent SDS, as assayed by colony forming units (CFU) 24 hours after treatment (49). When we examined the effect of Δ*sanA* on SDS sensitivity in carbon-limited stationary phase at shorter time scales, we found that the decrease in CFU began within the first hour of treatment and continued through 7 hours (**Figure 1A**). In contrast, SDS-treated wild-type cells maintained viability similar to that of untreated cells throughout the 7-hour treatment. An insertion mutation in *sanA* was reported to increase vancomycin sensitivity at high temperatures (43 °C) (58). To confirm this phenotype in a Δ*sanA* mutant, we performed efficiency-of-plating assays (EOPs) with wild-type and Δ*sanA* cells carrying either an empty vector or a low copy number plasmid expressing *sanA*. When treated with vancomycin at 43 °C, we found the Δ*sanA* strain was more sensitive to vancomycin and this phenotype could be complemented (**Figure 1B**). To confirm if we observed a true change in resistance (i.e., the concentration of antibiotic that can kill cells), we performed minimum inhibitory concentration (MIC) assays and found that the MIC of vancomycin for the Δ*sanA* strain was significantly lower than that of wild type (**Figure 1C**).

**Figure 1:**
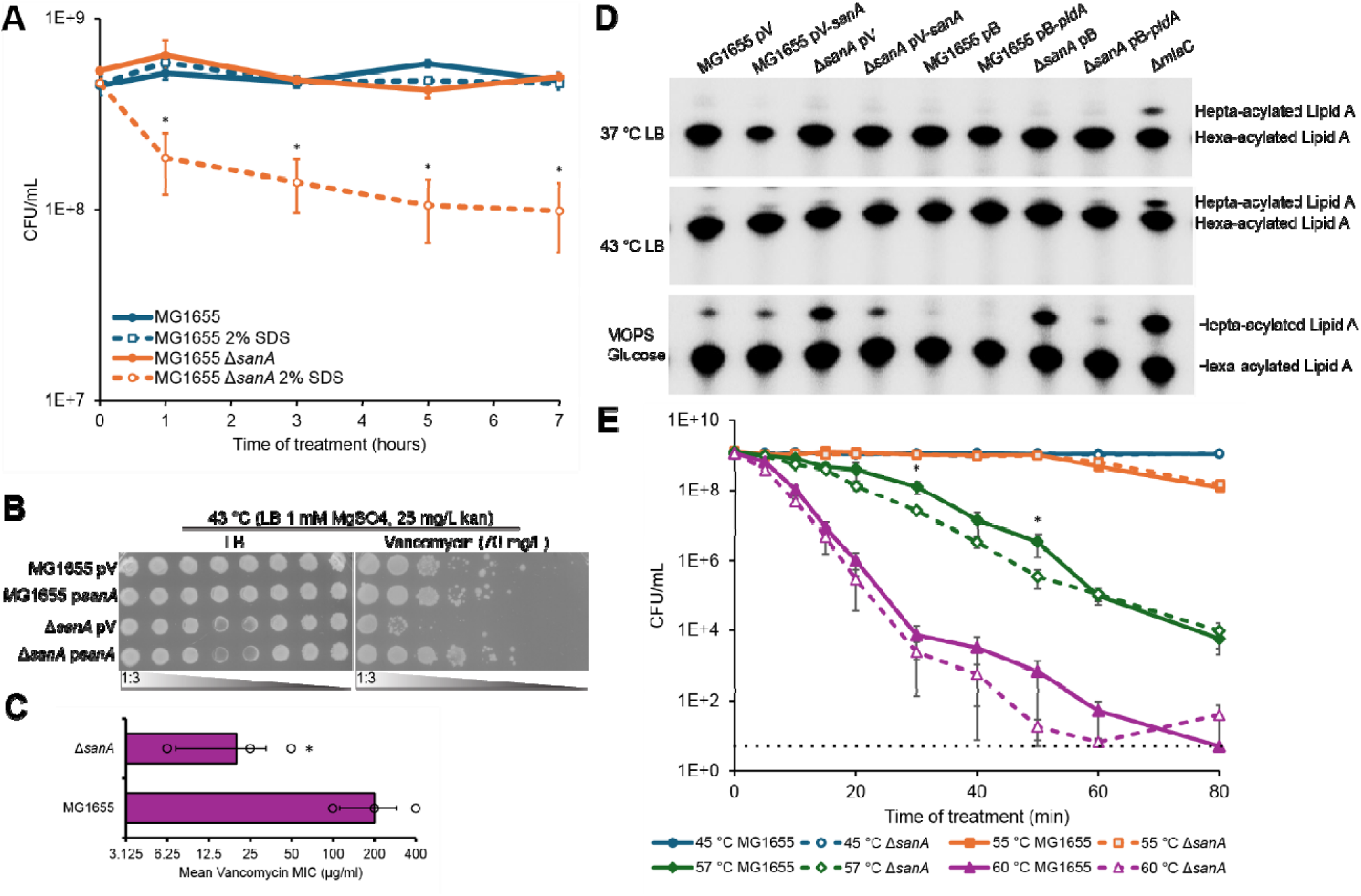
Loss of SanA causes increased envelope permeability and low levels of temperature sensitivity. **(A)** The indicated strains were grown to carbon-limited stationary phase and then treated with SDS-EDTA or a vehicle control and viability was assayed by CFU. The Δ*sanA* strain is significantly more sensitive to SDS than the wild-type strain from early time points post treatment. **(B)** The sensitivity of the indicated strains to vancomycin when grown at 43 °C was assayed by EOP. Plates were supplemented with MgSO_4_ to support growth. The Δ*sanA* strain was more sensitive to vancomycin than the wild-type strain and this sensitivity could be complemented with plasmid-based expression of *sanA*. **(C)** To confirm a true change in vancomycin resistance, the MIC for vancomycin was assayed for the wild-type and Δ*sanA* strain. The Δ*sanA* strain was significantly more sensitive to vancomycin than the wild-type strain. **(D)** Modifications to LPS were assayed by thin layer chromatography (TLC) for strains grown in the indicated conditions. The bands for hexa- and hepta-acylated lipid A are shown. The Δ*sanA* strain shows an increase in hepta-acylated lipid A which can be mitigated by complementation of *sanA* or overexpression of *pldA*. The *mlaC* strain is known to have an increase in phospholipids in the outer leaflet of the OM (68) leading to increased helpta-acylated lipid A and serves as a control. **(E)** Cells were grown to carbon-limited stationary phase at 37 °C and then their ability to survive at the indicated temperatures was assayed. The Δ*sanA* strain shows a significant decrease in survival at 57 °C. Quantitative data are the average of three biological replicates ± the SEM. For MICs, data are shown as the geometric average with individual data points. Images are representative of three independent experiments. * p<0.05 by the Mann Whitney Test.

Vancomycin is a large glycopeptide antibiotic for which entry into the cell is greatly impeded by LPS in the OM (4, 63, 64) and anionic detergents such as SDS are excluded from the cell by the strong hydrophilic network of lateral interactions between LPS molecules in the OM (1, 4). Given the importance of LPS for exclusion of these molecules, we examined the Δ*sanA* strain under various growth conditions for possible changes in the lipid A component of LPS. Lipid A anchors LPS in the OM and is critical for maintaining the OM permeability barrier. We observed a large increase in hepta-acylated lipid A in the Δ*sanA* mutant in cells grown in defined MOPS media with glucose but not in cells grown in LB media at 37 or 43 °C (**Figure 1D, S1**). The hepta-acylated lipid A levels were returned to normal by plasmid-based complementation of *sanA*. Hepta-acylated LPS is formed when PagP, an OMP, transfers a saturated acyl chain to LPS from a phospholipid mislocalized to the outer leaflet of the OM (65). Thus, increased hepta-acylated LPS indicates a loss of OM asymmetry causing increased phospholipids on the cell surface (66). In support of this loss of asymmetry, overexpression of *pldA*, an OM phospholipase that degrades phospholipids mislocalized to the OM (67, 68), decreased the amount of hepta-acylated lipid A in the Δ*sanA* strain (**Figure 1D, S1**). However, *pldA* overexpression did not restore the Δ*sanA* mutant’s vancomycin resistance at 43 °C or SDS resistance during carbon limitation (**Figure S2**), suggesting the loss of OM asymmetry was secondary to another defect that caused OM permeability.

SanA only affects vancomycin resistance at high temperatures (58) and *sanA* is predicted to have a σ^H^ promoter (56, 57). Therefore, we investigated whether the loss of SanA affected heat shock resistance in carbon-limiting conditions. Compared to the wild-type strain, the Δ*sanA* strain showed a small but significant decrease in survival at 57 °C (**Figure 1E**). This increased sensitivity was not shared with deletion of the other gene in *sanA*’s operon, *yeiS* (**Figure S3**). Overall, these data demonstrate that SanA plays an important role in maintaining the OM permeability barrier specifically in conditions of cellular stress such as carbon limitation and high temperatures.

### *sanA* has genetic interactions with ECA mutations that induce PG stress

To investigate how SanA affects membrane permeability, we made double mutants of *sanA* and non-essential members of various envelope biosynthesis pathways. A paralog of *sanA*, *elyC*, has been shown to have genetic interactions with ECA mutations (69) and is involved in regulating the synthesis of one form of ECA (70). In addition, in the same screen as *sanA*, we identified another gene, *yhdP*, which encodes a putative intermembrane phospholipid transporter (50–53), and *yhdP*’s membrane permeability phenotypes are dependent on the presence of ECA (49). Therefore, we decided to assay genetic interactions between ECA biosynthesis genes and *sanA*.

We found that Δ*sanA* had a synthetic SDS-EDTA sensitivity, even at 37 °C in LB media, when paired with a deletion of *wecA*, the first gene in the ECA synthesis pathway (**Figure 2A**). EDTA breaks the network of interactions between LPS molecules, increasing cells susceptibility to SDS (71). Intriguingly, when we tested for genetic interactions between *sanA* and *wecE*, which encodes a protein involved in synthesis of the third sugar in the ECA trisaccharide, we found that Δ*wecE* causes SDS-EDTA sensitivity and this sensitivity is suppressed by Δ*sanA* (**Figure 2A**). These genetic interactions were apparent at 43 °C as well as 37 °C (**Figure S4A**) and the synthetic SDS-EDTA phenotype of the Δ*sanA* Δ*wecA* mutant could be complemented with plasmid-based expression of *sanA* (**Figure S4B**). The difference in phenotype between Δ*wecA* and Δ*wecE* suggested that interactions between ECA and other biosynthetic pathways are important for this phenotype, not the loss of ECA itself. Specifically, ECA synthesis uses the same isoprenoid carrier (Und-P) as PG synthesis (33). Therefore, we wondered if PG synthesis stress caused by sequestration of the isoprenoid carrier might be important for the genetic interaction with *sanA*.

**Figure 2:**
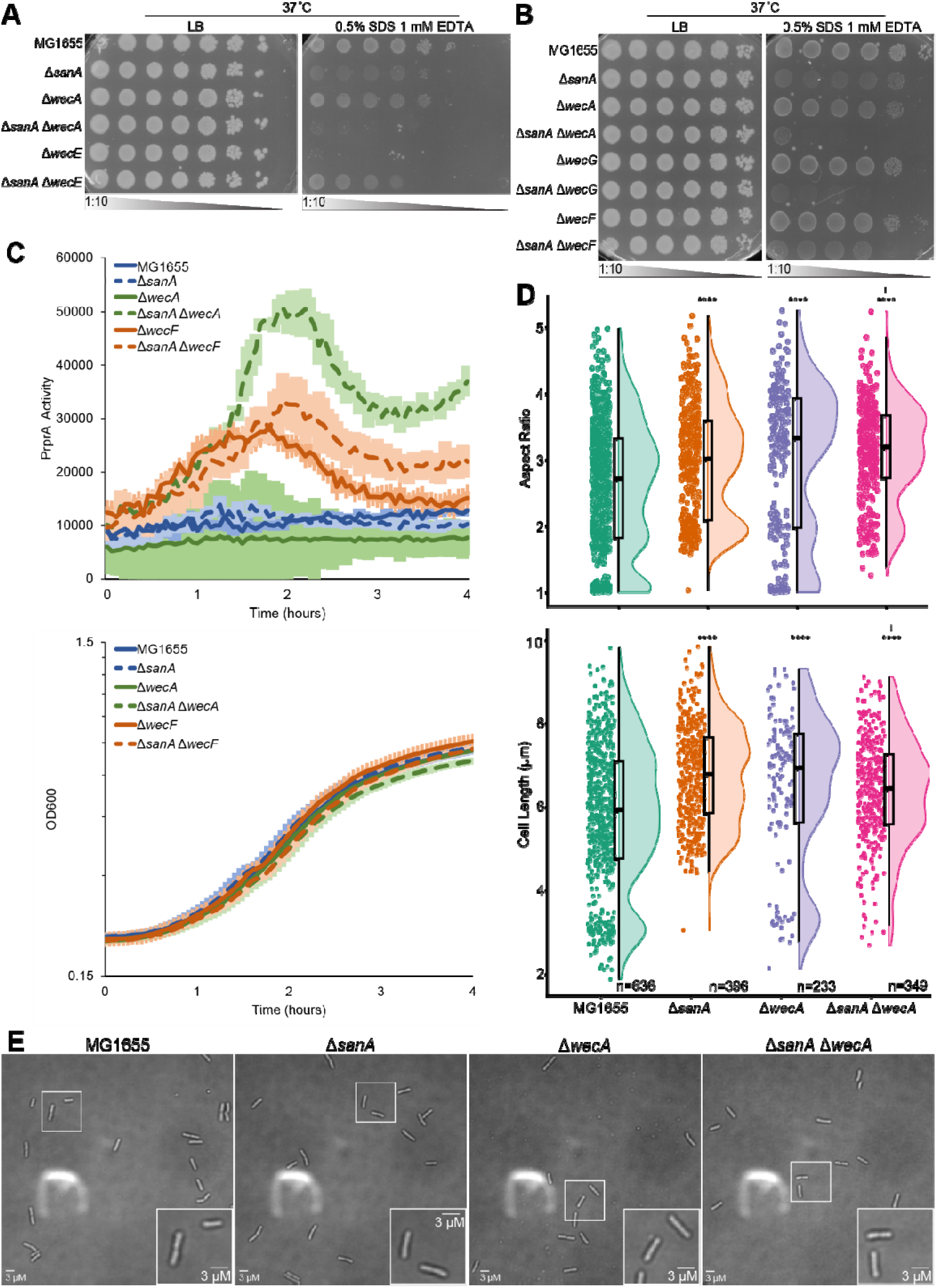
SanA shows genetic interactions with ECA synthesis mutations differentially affecting PG stress. **(A-B)** The indicated strains were assayed for SDS-EDTA resistance by EOP. Images are representative of three independent experiments. (A) A double mutant of Δ*sanA* Δ*wecA* showed synthetic SDS-EDTA sensitivity, while Δ*sanA* suppressed the SDS-EDTA sensitivity of the Δ*wecE* strain. (B) Double mutants of each ECA transglycosylase showed less SDS-EDTA sensitivity than the previous strain. **(C)** Activation of the Rcs stress response in the indicated strains was assayed using a *luxCDABE* reporter. Data are shown as luminescence relative to OD_600_. The growth of the cultures (OD_600_) over the course of the experiment is also shown. Data are the average of three biological replicates ± the SEM. **(D)** Cell shape of the indicated strains was assayed, and the length and aspect ratio (length/width) were calculated. Raincloud plots are shown with individual data points, the population distribution, and a box plot. **** p<0.00005 vs. wild type MG1655 by Wilcoxon Test; ‡ p<0.05 vs. the Δ*sanA* strain by Wilcoxon Test. **(E)** Representative images demonstrating the cell shape of the wild type Δ*sanA*, Δ*wecA*, and Δ*sanA* Δ*wecA* strains. Insets display the indicated regions.

We proceeded to compare the phenotypes of Δ*sanA* with deletion of the three transglycosylases in the ECA synthesis pathway. The PG stress caused by deletion of each subsequent transglycosylase is more pronounced than the previous (37, 40, 69, 72). Δ*wecA* does not cause apparent PG stress, Δ*wecG* causes minimal PG stress, and Δ*wecF* causes substantial PG stress as evidenced by envelope stress response activation, permeability defects, and cell shape defects, although these defects are not as strong as those of Δ*wecE* (37, 40, 69, 72). When we tested these mutants, we indeed found the Δ*sanA* Δ*wecA* strain to have the highest SDS-EDTA sensitivity, while the Δ*sanA* Δ*wecG* strain had slight sensitivity and the Δ*sanA* Δ*wecF* strain showed similar SDS-EDTA sensitivity to that of the Δ*sanA* strain (**Figure 2B**). These data indicated that, counterintuitively, the envelope permeability of a Δ*sanA* mutant is increased by mutations that increase isoprenoid carrier availability.

ECA mutants causing PG stress have been shown to activate the Rcs stress response, which directly responds to LPS defects but can also be activated by PG stress (73–75). Therefore, we tested Rcs activation using luciferase genes driven by the *rprA* promoter (76, 77) in single and double deletion mutants of *sanA*, *wecA* and *wecF*. Neither Δ*sanA* or Δ*wecA* alone caused Rcs activation (**Figure 2C**); however, a Δ*sanA* Δ*wecA* mutant showed Rcs activation greater than that of the Δ*wecF* or Δ*sanA* Δ*wecF* mutant. The peak reporter activity in the Δ*sanA* Δ*wecA* strain corresponded to exponential growth. As these data and the SDS-EDTA data point to interactions with isoprenoid carrier availability and PG synthesis, we then assayed the cell shape of the Δ*sanA* and Δ*wecA* single and double mutants. We grew cells for 7-10 generations in exponential phase before the assay to ensure that cell shape was stable after exiting stationary phase (78, 79). The aspect ratio (length/width) of all mutants was larger than wild type with the Δ*sanA* Δ*wecA* mutant showing a significant shape change from that of the Δ*sanA* single mutant (**Figure 2DE**). Overall, these data demonstrate *sanA* has genetic interactions causing increased permeability and envelope stress with increased isoprenoid carrier availability.

### Spontaneous mutation to *ftsI* and *prc* suppress the Δ*sanA* Δ*wecA* SDS-EDTA sensitivity

When assaying the SDS-EDTA sensitivity of the Δ*sanA* Δ*wecA* double mutant, we occasionally noticed the appearance of spontaneous suppressor mutants. Therefore, we decided to select for these mutants to determine what types of mutations could suppress this strain’s envelope permeability. We plated Δ*sanA* Δ*wecA* strain on a concentration of SDS-EDTA where it did not grow but the wild-type strain did and collected 11 suppressor mutants from a total of 7 separate cultures. We grouped these mutants by their antibiotic resistance patterns to find strains with similar mutations (**Figure S5**) and then performed whole genome sequencing on representatives of each group, following up to confirm the other mutants within the group had the same mutation(s). These suppressor mutants each contained one of four mutations, *ftsI_V98E_*, *ftsI_S165P_*, *prc_F365L_*, or *prc_R578C_* (**Figure 3A**). FtsI is the DD-transpeptidase involved in septal PG synthesis during cell division (41, 80), while Prc (a.k.a. Tsp) is a protease known to cleave FtsI’s C-terminal 11 amino acids to produce the fully mature protein (61, 62). As Prc is necessary for FtsI maturation (61), we focused on the *ftsI* mutations as the more direct suppressors. However, it is possible that the *prc* mutations may act through an unrelated pathway (i.e., by increasing endopeptidase levels, which has been recently shown to suppress another Δ*sanA* phenotype) (24, 81). We linked the *ftsI_V98E_* mutation to a nearby Tn*10* and used this marker to transfer the mutation to the wild-type, Δ*sanA*, Δ*wecA*, and Δ*sanA* Δ*wecA* backgrounds. After being moved to a clean background, the *ftsI_V98E_* mutation suppressed SDS-EDTA sensitivity in the Δ*sanA* Δ*wecA* strain showing it was sufficient for suppression (**Figure 3B**). We were not successful in moving the *ftsI_S165P_* mutation to other strain backgrounds; however, we used P1*vir* transduction to remove the mutation from the suppressor strain and this prevented the SDS-EDTA resistance of the strain (**Figure 3C**). Although we assayed the effect of the *ftsI_V98E_* mutation on the vancomycin sensitivity of a Δ*sanA* strain at 43 °C, the *ftsI_V98E_* mutation also caused vancomycin resistance in a wild-type background making it difficult to assess whether its effect on the Δ*sanA* vancomycin sensitivity was specific (**Figure S6**).

**Figure 3:**
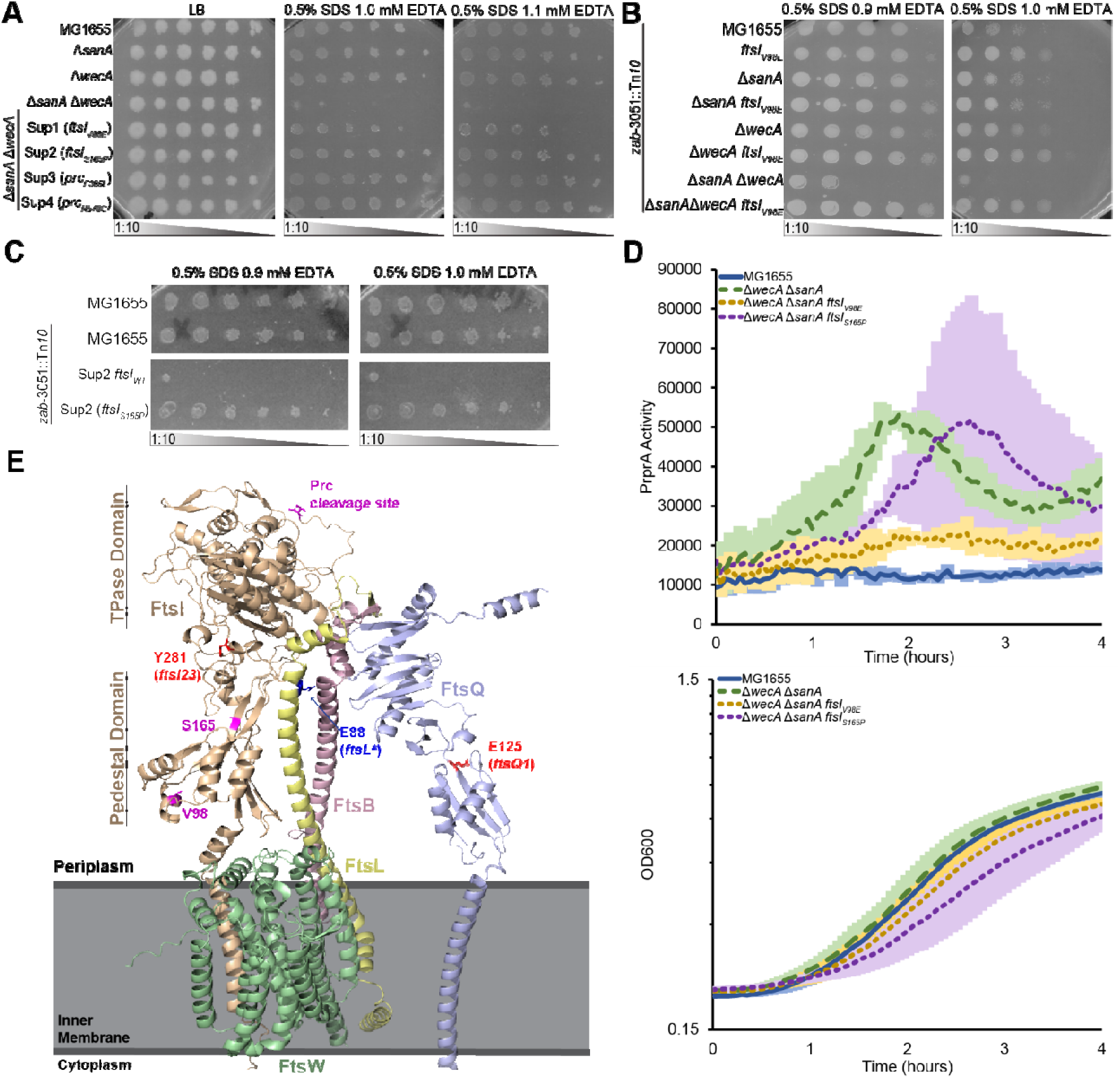
The SDS-EDTA of a Δ*sanA* Δ*wecA* strain can be suppressed through mutations to *ftsI* and *prc*. **(A-B)** The SDS-EDTA sensitivity of the indicated strains was assayed by EOP. Images are representative of three independent experiments. (A) Suppressor mutations show increased SDS-EDTA resistance compared to the parent Δ*sanA* Δ*wecA* strain. (B) A linked Tn*10* was used to transfer the *ftsI_V98E_* mutation the indicated strains. The *ftsI_V98E_* mutation was sufficient for suppression. (C) The *ftsI_S165P_* mutation was removed from the indicated suppressor and the loss of this mutation prevented suppression demonstrating that it is necessary for suppression. **(D)** Reporter assays for Rcs activation were performed as in Figure 2C. The P*_rprA_* activity (luminescence/OD_600_) as well as the growth of the strains over time are shown. The *ftsI_V98E_* mutation suppressed the Rcs activation phenotype of the Δ*sanA* Δ*wecA* strain but the *ftsI_S165P_* mutation did not. The Δ*sanA* Δ*wecA* strain with the *ftsI_S165P_*mutation also had a noticeable growth defect. Data are the average of three biological replicates ± the SEM. **(E)** A cartoon of the FtsQLBIW complex modeled with AlphaFold 3 is shown (82). V98 and S165, as well as the localization of other selected variants, are shown in stick format. The locations of the FtsI periplasmic domains, the pedestal domain and the transpeptidase (TPase) domain are indicated.

We then asked whether the *ftsI* mutations could suppress Rcs activation in the Δ*sanA* Δ*wecA* strain. We found that the *ftsI_S165P_* mutation caused a growth delay and maintained Rcs activation, while the *ftsI_V98E_* mutation brought Rcs activation in the Δ*sanA* Δ*wecA* mutant back to nearly wild-type levels without a significant effect on growth (**Figure 3D**). We made an AlphaFold model (82) of a partial divisome containing FtsI, FtsW, FtsL, FtsB, and FtsQ, which corresponds well to the crystal structure of this complex in *Pseudomonas aeruginosa* (83), and mapped locations of the suppressor mutations onto this predicted structure (**Figure 3E**). FtsI has two periplasmic domains, a pedestal domain thought to control activation of FtsW and mediate interactions with FtsL and FtsB and the catalytic transpeptidase domain (80, 84). The Prc cleavage site in FtsI is in an unstructured region adjacent to the transpeptidase domain, while the V98E and S165P mutations are in the pedestal domain (**Figure 3E**). Neither of these mutations has been previously reported and their effect on FtsI activity is yet to be determined. Nevertheless, the consistent and repeated isolation of mutations in *ftsI* and *prc* demonstrated that the envelope defect leading to SDS-EDTA sensitivity in the Δ*sanA* Δ*wecA* strain is related to cell division.

### Suppressor mutations in *ftsI* cause partial loss of FtsI function

As the Δ*sanA* and Δ*sanA* Δ*wecA* strains showed a cell shape with a higher aspect ratio than wild-type cells (**Figure 2D**) and the suppressor mutations we found are in cell division genes, we then assayed the cell shape of the suppressor strains. Cells from both *ftsI* mutant strains were significantly longer than the Δ*sanA* Δ*wecA* parent and had a greater aspect ratio (**Figure 4AB**). In contrast, the aspect ratio of the *prc* mutants was indistinguishable from that of the Δ*sanA* Δ*wecA* parent. The longer length of the *ftsI* mutants suggests that they have a partial loss of *ftsI* function. Therefore, we tested the transpeptidase activity of the mutants through resistance to aztreonam, a β-lactam antibiotic that is highly specific to FtsI (85).

**Figure 4:**
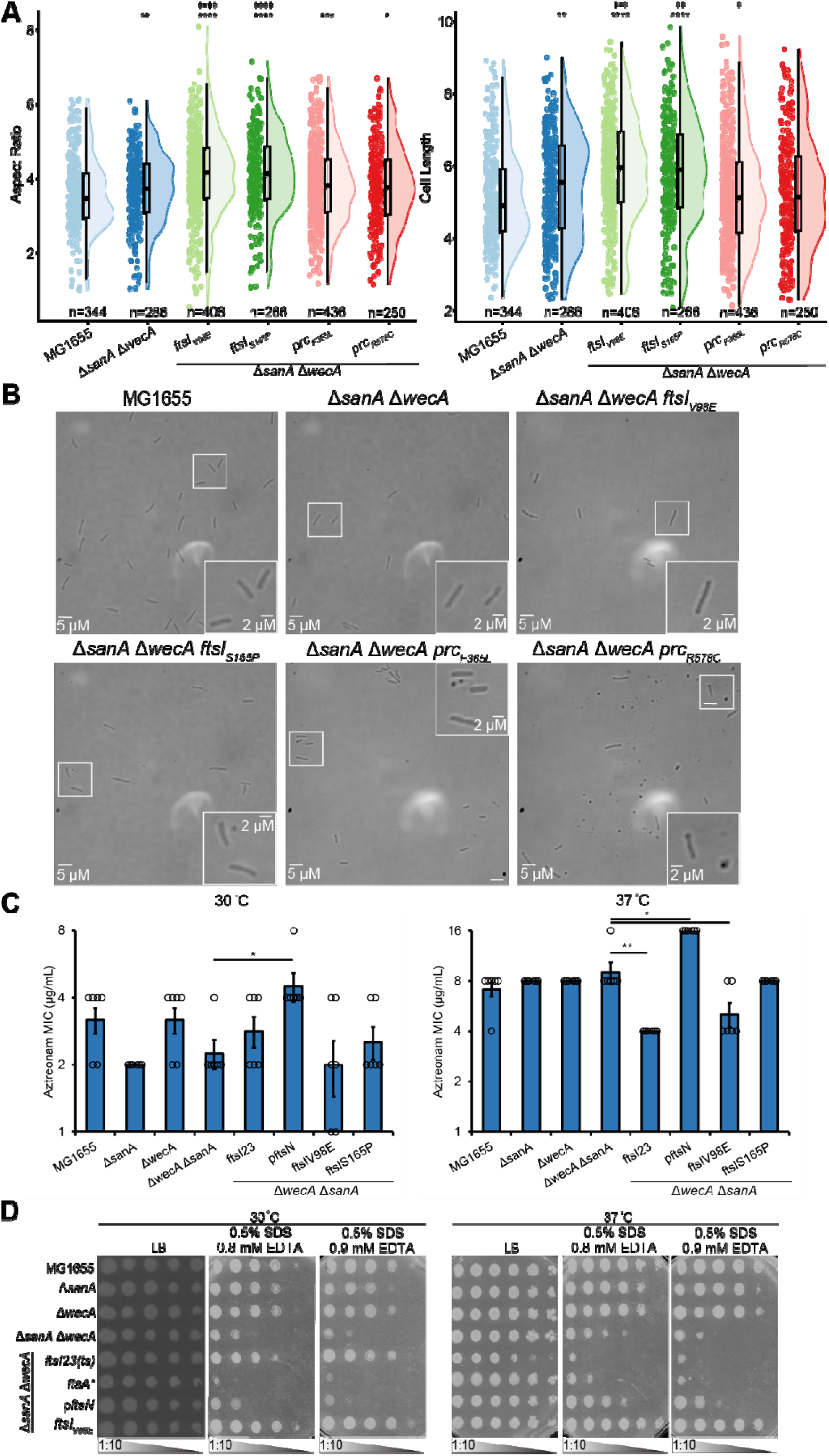
Suppressors of Δ*sanA* Δ*wecA* SDS-EDTA sensitivity cause partial loss of FtsI function. **(A)** Cell shape of the indicated strains was assayed, and the aspect ratio (length/width) and cell length are shown. Raincloud plots are shown with individual data points, the population distribution, and a box plot. * p<0.05, **p<0.005, ***p<0.0005, **** p<0.00005 vs. wild type MG1655 by Wilcoxon Test; ‡ p<0.05, ‡‡ p<0.005, ‡‡‡ p<0.0005, ‡‡‡‡ p<0.00005 vs. the Δ*sanA* Δ*wecA* strain by Wilcoxon Test. Mutations in *ftsI* caused the Δ*sanA* Δ*wecA* cells to become significantly longer, while *prc* mutations had less of an effect on length. **(B)** Representative images demonstrating cell shape are shown for the indicated strains. Insets display the indicated regions. **(C)** The sensitivity of the indicated strains to aztreonam, a β-lactam targeting FtsI, was assayed by MIC. Data are shown as the geometric average of six biological replicates ± the SEM with individual data points. * p<0.05, ** p<0.005 by Mann-Whitney Test. *ftsI23* and overexpression of *ftsN* serve as controls for decreased and increased activity of FtsI, respectively. The *ftsI_V98E_* mutation caused a significant increase in aztreonam sensitivity at 37 °C, indicating decreased FtsI transpeptidase activity. **(D)** The SDS-EDTA sensitivity of the indicated strains was assayed by EOP at both 30 and 37 °C. Of the cell division mutants, only *ftsI23* was able to suppress the SDS-EDTA sensitivity of the Δ*sanA* Δ*wecA* strain. Images are representative of three independent experiments.

To show the correlation between FtsI activity and aztreonam resistance, we tested *ftsI23* (Y281D), a temperature sensitive *ftsI* allele that has reduced activity even at permissive temperatures and can slow the rate of closure of the septum (86–88), and *ftsN* overexpression which has been shown to result in increased divisome activity and to suppress partial loss of function mutations in both divisome assembly and septal PG synthesis (89–91). Given the temperature sensitive phenotype of *ftsI23,* we assayed aztreonam MICs at both 30 and 37 °C. At 30 °C, *ftsN* overexpression caused a significant increase in aztreonam MIC in the Δ*sanA* Δ*wecA* background (**Figure 4C**), while none of the *ftsI* mutations caused a significant change in MIC. However, at 37 °C, *ftsI23* and *ftsN* overexpression caused a significant decrease and increase in MIC, respectively (**Figure 4C**). Thus, FtsI activity positively correlates to aztreonam resistance. At 37 °C, *ftsI_V98E_* decreased aztreonam’s MIC compared to its Δ*sanA* Δ*wecA* parent, while *ftsI_S165P_* did not cause a significant change. These data suggest that, while *ftsI_V98E_* may have a direct or indirect decreased transpeptidase activity contributing to increased cell length, *ftsI_S165P_*does not have a significant effect on transpeptidase activity. Therefore, the effect of these mutations on protein-protein interactions, rather than transpeptidase activity, may be more important for suppression.

To test if suppression of Δ*sanA* Δ*wecA* membrane permeability was unique to *ftsI* mutations, we tested the effect of several gain- and loss-of-function mutations in divisome activity on the SDS-EDTA sensitivity of the Δ*sanA* Δ*wecA* strain. In contrast to *ftsI_V98E_* that suppressed the SDS-EDTA sensitivity of the Δ*sanA* Δ*wecA* strain at both 30 and 37 °C (**Figure 4D**), *ftsI23* suppressed only at 30 °C, suggesting there may be a range of function loss for *ftsI* that can suppress envelope permeability. Neither *ftsA\** (R286W) (92, 93) nor *ftsN* overexpression, both of which are activating mutations, suppressed the SDS-EDTA sensitivity of the Δ*sanA* Δ*wecA* mutant (**Figure 4D**). In fact, the Δ*sanA* Δ*wecA* strain carrying the *ftsA\** allele was more SDS-EDTA sensitive than Δ*sanA* Δ*wecA* parent. Overall, these data demonstrate that other loss of function mutations in *ftsI* can also suppress the Δ*sanA* Δ*wecA* envelope permeability.

### Δ*sanA* Δ*wecA* envelope permeability is linked to septal PG synthesis

To test whether SanA might influence the timing of Z ring assembly, the first step in cell division, we expressed GFP-labeled FtsZ (94) and failed to observe significant differences in the localization of FtsZ in the wild-type, Δ*sanA*, and Δ*sanA ftsI_V98E_* strains (**Figure S7AB**). Moreover, the *ftsI_V98E_* mutation did not change FtsN localization in a Δ*sanA* Δ*wecA* background (**Figure S7CD**). Therefore, we examined other cell division mutations to determine which components of the divisome, its assembly and/or septal PG synthesis, were important for suppression of SDS-EDTA sensitivity in the Δ*sanA* Δ*wecA* strain. We examined the effect of three temperature sensitive loss-of-function mutations in divisome assembly *ftsZ84* (G105S) (95), *ftsA12* (A188V) (96, 97), and *ftsA27* (S195P) (90, 98), as well as a gain-of-function mutation, *ftsL\** (E88K) (99, 100), and a temperature sensitive loss-of-function mutation in septal PG synthesis, *ftsQ1* (E125K) (96, 101). Given the temperature sensitivity caused by these mutations we performed the experiment at 30 °C. None of the mutations affected SDS-EDTA sensitivity on their own except *ftsQ1*, which showed sensitivity similar to that of the Δ*sanA* Δ*wecA* strain (**Figure 5A**). Interestingly, only *ftsQ1* suppressed the SDS-EDTA sensitivity of the Δ*sanA* Δ*wecA* strain (**Figure 5B**), demonstrating cross suppression between these mutations. In addition, we found that the *ftsZ84*, *ftsA12*, and *ftsL\** mutations cause SDS-EDTA sensitivity in the Δ*sanA* background (**Figure 5C**). Overall, these data demonstrate that the increased envelope permeability of the Δ*sanA* Δ*wecA* strain is suppressed by septal PG synthesis mutants but not by mutations in divisome assembly. Instead, divisome assembly mutants can increase the envelope permeability of a Δ*sanA* strain.

**Figure 5:**
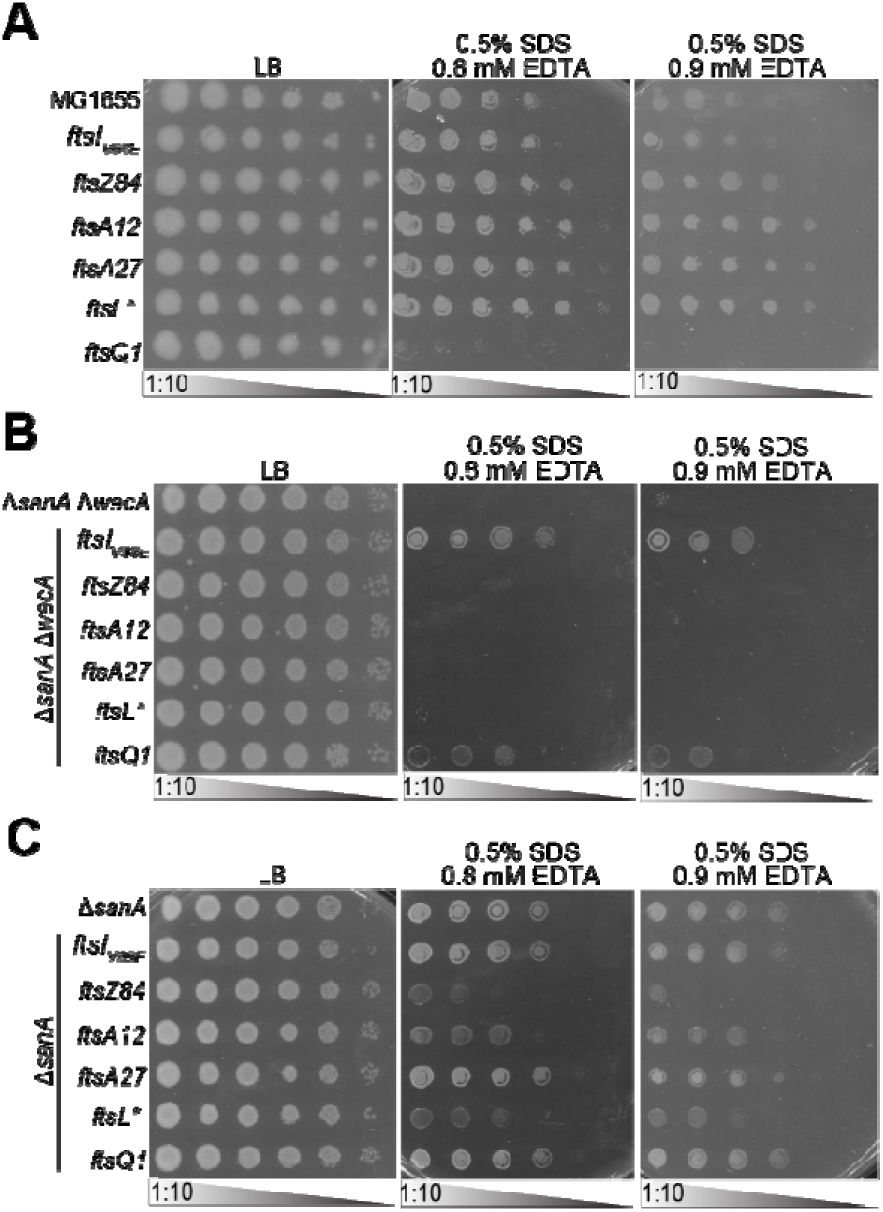
Suppression of Δ*sanA* Δ*wecA* phenotypes is linked to septal PG synthesis and not divisome assembly. EOPs were performed at 30 °C to assay the SDS-EDTA sensitivity of the indicated strains. The effect of cell division mutations in a wild-type background (A), a Δ*sanA* Δ*wecA* background (B), and a Δ*sanA* background (C) are shown. Images are representative of three independent experiments. **(A)** Only the *ftsQ1* mutation causes SDS-EDTA sensitivity in a wild-type background. **(B)** Of the assayed mutations, only *ftsQ1* suppressed the SDS-EDTA sensitivity of the Δ*sanA* Δ*wecA* strain, while earlier loss-of-function mutations in divisome assembly or gain-of-function mutations in septal PG synthesis did not. **(C)** The *ftsZ84, ftsA12,* and *ftsL\** mutations make the Δ*sanA* strain more sensitive to SDS-EDTA.

To directly evaluate the effect of *sanA* and *ftsI_V98E_* on septal PG synthesis, we incubated cells with BODIPY-FL-3-amino-D-alanine (BADA) for 10 minutes to label newly synthesized PG and compared PG incorporation at center-cell regions (center line ±5% of cell length) vs. the whole cell in dividing and non-dividing cells (**Figure 6**). In both dividing and non-dividing cells, Δ*sanA* mutants exhibited a lower ratio of center-localized to total PG incorporation than the wild type, suggesting a spatial redistribution of PG synthesis. In non-dividing cells, there was no difference between the Δ*sanA* and Δ*sanA ftsI_V98E_* strain. However, in dividing Δ*sanA ftsI_V98E_* cells, center cell PG synthesis was similar to that of wild type and significantly greater than that of Δ*sanA*. These data suggest that septal PG synthesis is increased with the *ftsI* mutation, which is an interesting contrast to its longer cell length.

**Figure 6:**
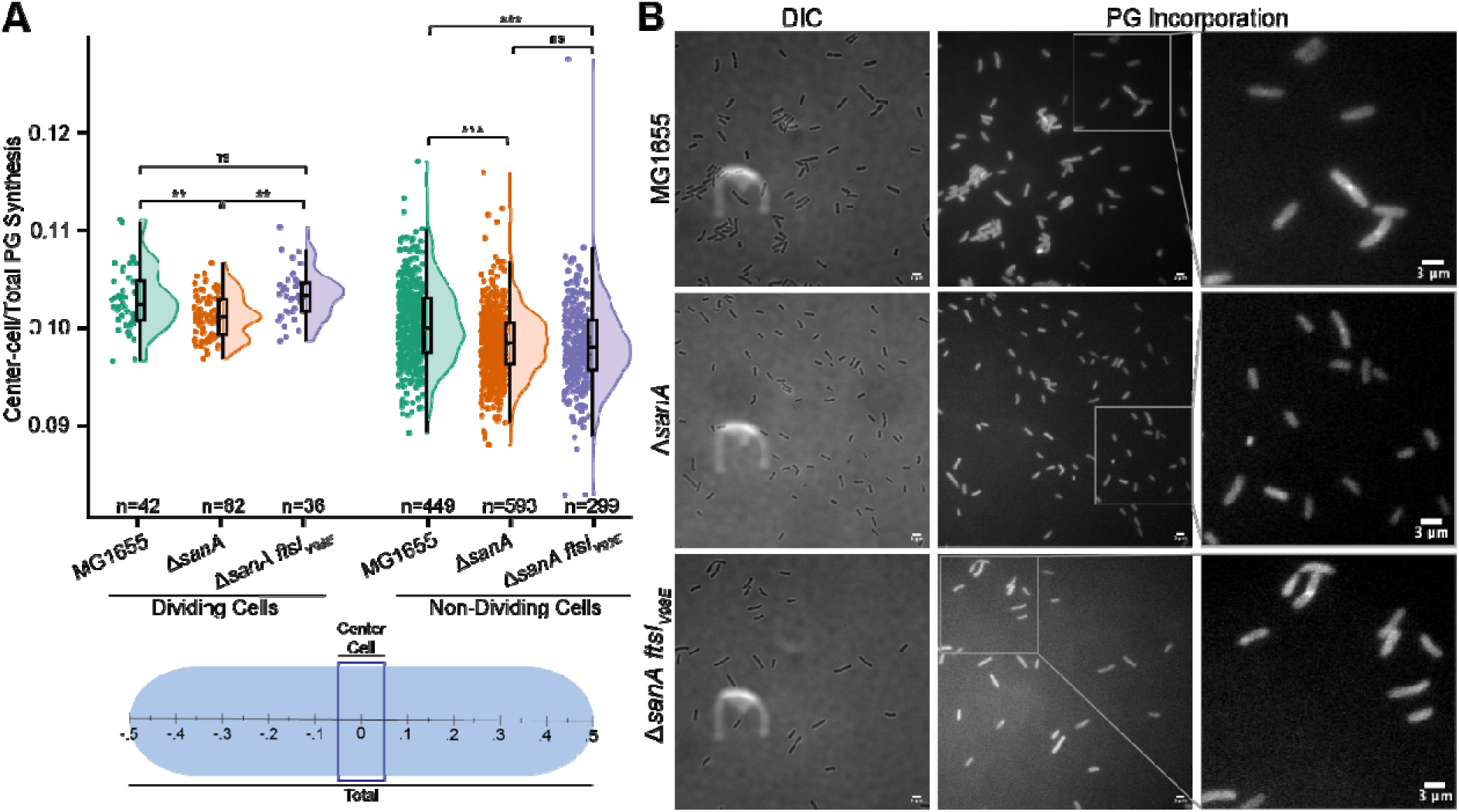
FtsI suppressor restores balance between septal and elongation PG synthesis in a Δ*sanA* strain. **(A)** To compare septal PG and elongation PG synthesis, newly synthesized PG was labeled with fluorescent D-ala (BADA) and cells were imaged. The ratio of new PG incorporation at the center 10% of the cell versus the total cell was calculated. In both dividing and non-dividing cells, this ratio was lower in Δ*sanA* than wild-type cells. However, the *ftsI_V98E_* suppressor in the Δ*sanA* background restored the ratio to wild type levels only in dividing cells. Representative DIC and fluorescence images showing PG incorporation are shown for the indicated strains. Expanded regions of fluorescence images are taken from the indicated regions. Scale bars are 3 µm.

### Membrane permeability with loss of SanA is tied to lipid II availability and FtsW activity

As the *ftsI* suppressor mutations reside in the FtsI pedestal domain that interacts with FtsW and the *ftsI_V98E_* mutation increases septal PG synthesis, we investigated whether SanA’s envelope permeability phenotypes relate to FtsW activity. We transferred a previously described (102) FtsW depletion system that contains an arabinose-inducible *ftsW* expressing plasmid and chromosomal Δ*ftsW* to our strains. We found that *ftsI_V98E_*could suppress the Δ*sanA* Δ*wecA* strain regardless of arabinose concentration (**Figure 7A**). However, *ftsI_V98E_* in a wild-type background conferred resistance to arabinose depletion, allowing cells to grow at a lower concentration of arabinose than is tolerated by the parent strain, suggesting increased FtsW activity (**Figure 7A**). We did not observe the same increase when *ftsI_V98E_* is expressed in either the Δ*sanA* or Δ*sanA* Δ*wecA* background, suggesting that loss of SanA may increase the activity of FtsW necessary for viability.

**Figure 7:**
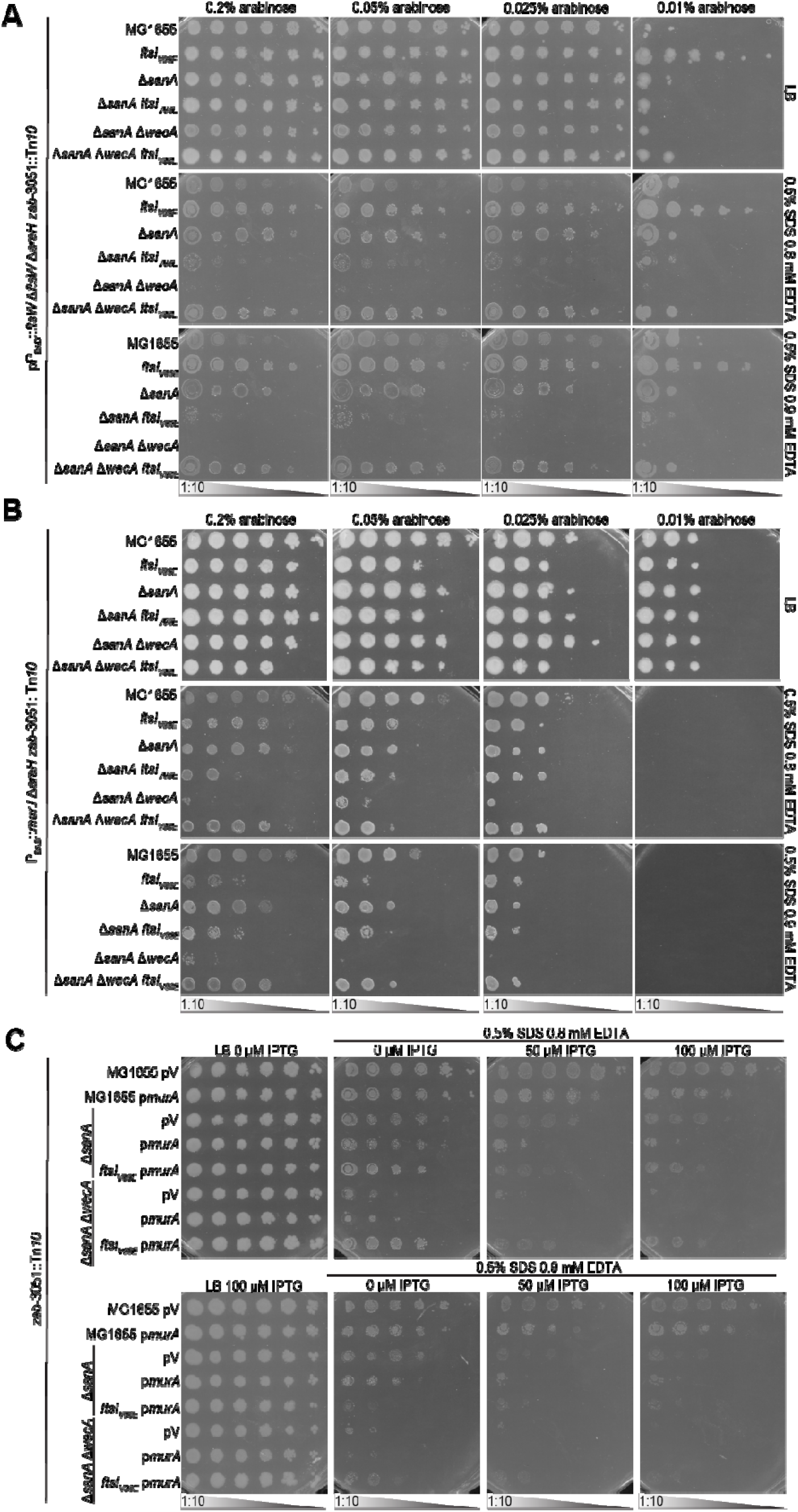
SanA membrane permeability phenotypes are tied to lipid II availability and FtsW activity. EOPs were performed to assay SDS-EDTA resistance in the following strains in the indicated conditions. Images are representative of three independent experiments. **(A)** FtsW depletion strains were constructed by deleting *ftsW* in strains carrying a plasmid-based copy of *ftsW* under an arabinose-inducible promoter. Deletion of *araH*, an inducible arabinose permease, ensures linear induction. Wild-type cells carrying the *ftsI_V98E_* mutation could grow at a lower arabinose concentration, indicating increased activity of FtsW. Interestingly, this increased activity is not observed with the *ftsI_V98E_* mutation in the Δ*sanA* or Δ*sanA* Δ*wecA* strains. **(B)** MurJ depletion constructs were made by transduction of an arabinose inducible promoter to replace the native *murJ* promoter. Depletion of MurJ did not prevent suppression of Δ*sanA* Δ*wecA* SDS-EDTA sensitivity by the *ftsI_V98E_*mutation. **(C)** An IPTG inducible *murA* overexpression plasmid or a vector control were introduced into the indicated strains. Overexpression of *murA* in a Δ*sanA* strain phenocopies the SDS-EDTA sensitivity of the Δ*sanA* Δ*wecA* strain and this sensitivity can be suppressed by the *ftsI_V98E_*mutation.

Given *sanA*’s genetic interaction with mutations altering isoprenoid carrier availability, we further investigated whether the availability of lipid II, the substrate for PG synthesis, was involved in SanA’s envelope permeability phenotype. A chromosomal depletion strain for MurJ, the lipid II flippase (102, 103), where the promoter for *murJ* was replaced with the arabinose-inducible P_BAD_ promoter has been described (103). We transduced this promoter into our strains and found MurJ depletion did not prevent suppression of Δ*sanA* Δ*wecA* SDS-EDTA sensitivity by *ftsI_V98E_* (**Figure 7B**). To test if increasing lipid II by overproducing MurA, which catalyzes the first committed step in PG synthesis (104), would affect OM permeability, we overexpressed *murA* in a Δ*sanA* strain. We observed SDS-EDTA sensitivity that phenocopied that observed in a Δ*sanA* Δ*wecA* strain and this sensitivity could be suppressed by *ftsI_V98E_*(**Figure 7C**). Furthermore, overexpression of *murA* decreased the suppression observed in a Δ*sanA* Δ*wecA ftsI_V98_* strain. These data demonstrate that multiple mutations expected to increase lipid II availability lead to SDS-EDTA sensitivity in the absence of SanA and that this sensitivity can be suppressed through mutations in septal PG synthesis genes.

### A predicted pocket is important for SanA’s function in envelope permeability

To further investigate how SanA’s structure affects its function, we modeled SanA’s structure using AlphaFold 3 (82). Using the predicted structure as a guide, we constructed point mutations to several conserved SanA residues (**Figure 8A**). Interestingly, SanA’s predicted structure has a pocket that faces the direction of the transmembrane helix and an amphipathic helix perpendicular to the transmembrane helix that may sit in the IM. In an attempt to validate this prediction, we first made charge switch mutation pairs of residues interacting with this amphipathic helix. Whereas two of the individual mutations (R152E and E182R) resulted in loss of *sanA* function (**Figure 8B**) without affecting protein stability (**Figure 8C**), a double mutation (R152E E182R) did not restore function. Mutations of another charge pair D192 and R198 resulted in loss of protein stability, complicating analysis (**Figure 8BC**). Therefore, we could not validate the interactions of the amphipathic helix. However, several structural models (AlphaFold 2, AlphaFold 3, and I-TASSER (82, 105, 106)) all predicted a pocket that contains several conserved residues (**Figure 8A**). Mutating conserved hydrophilic residues in the pocket resulted in loss of SanA function (**Figure 8D**), while generally maintaining protein stability (**Figure 8C**). Previous work also found the H149A mutation to cause loss of function (81). While charged residues are predicted near the outer edge of the pocket, the inner surface of the pocket is composed of hydrophobic residues (**Figure 8A**). Mutations to residues on the surface of the pocket (L51, G52) resulted in loss of function with minimal or no effect on protein stability, while mutations of buried residues did not change SanA function (**Figure 8CE**). Finally, we made several mutations to surface residues of SanA including those on the side of SanA predicted to face the IM and the side predicted to face away from the IM (**Figure 8A**). Generally, these mutations caused less severe phenotypes than mutations to SanA’s pocket residues (**Figure 8F**). Overall, these structure-function analyses suggest that SanA’s pocket may be important for binding or enzymatically altering an IM molecule.

**Figure 8:**
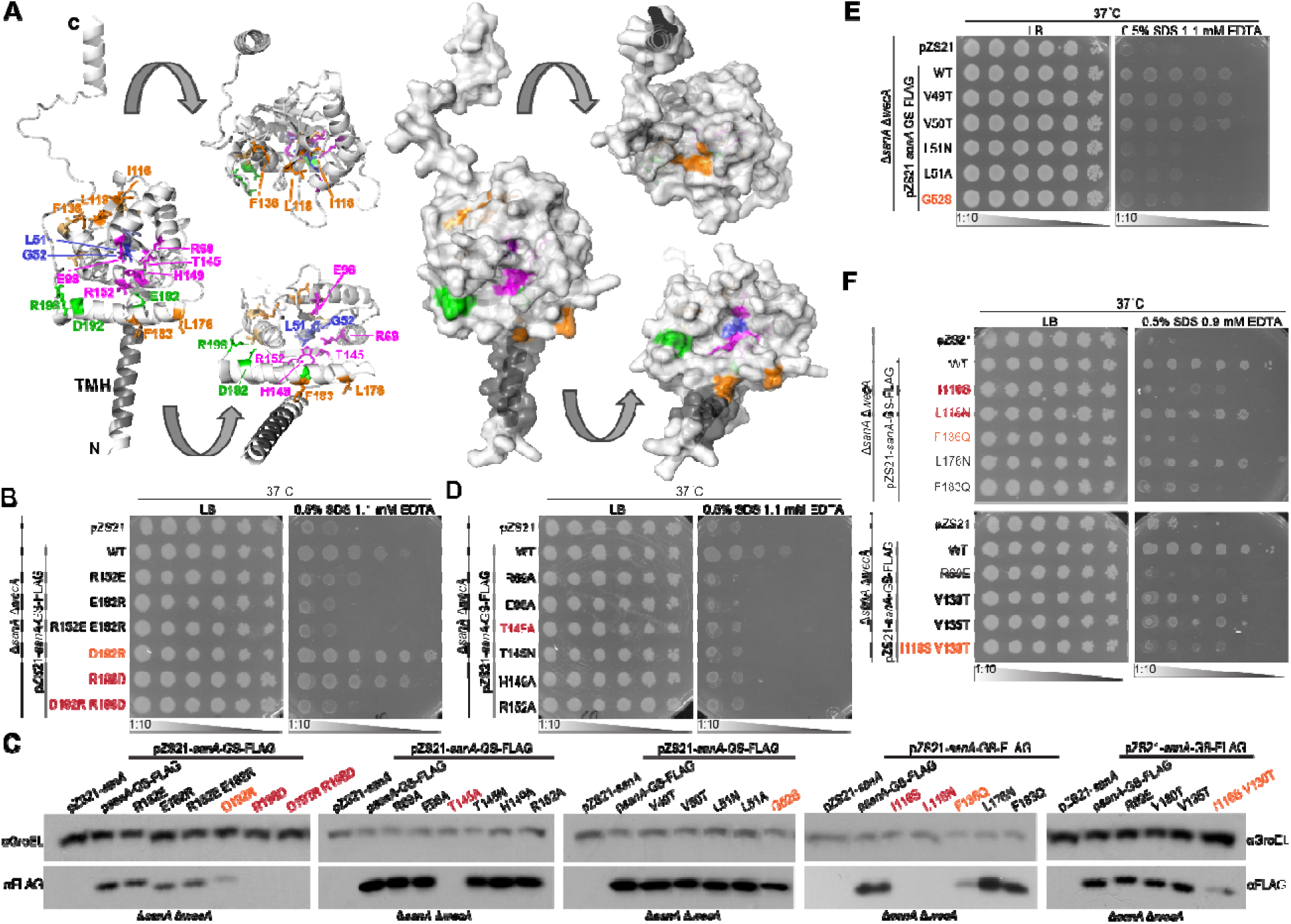
Predicted SanA pocket is important for SanA function. **(A)** EOPs were performed to assay SDS-EDTA resistance of the indicated strains. The effect of mutations of *sanA* charged residues predicted by AlphaFold 3 (82) to interact is shown. The mutations decrease SanA function and this function is not restored by double mutants expected to restore interaction. **(B)** Cartoon and surface predicted structures of SanA are shown with mutated residues labeled. Charge pair residues are shown in green, hydrophilic pocket residues in magenta, hydrophobic pocket residues in blue, and surface residues in orange. N: Cytoplasmic N-terminus; TMH: Transmembrane helix; C: C-terminus. **(C)** The stability of proteins produced by the C-terminally FLAG-tagged SanA mutants was assayed by immunoblot. Constructs with reduced protein accumulation are marked in orange and with undetectable protein accumulation in red. Images are representative of three independent experiments. GroEL serves as a loading control. **(D)** Mutations were made to conserved polar and charged residues in the predicted pocket of SanA and SanA function was assayed by EOP. These mutations impede SanA function. **(E)** Mutations were made to conserved non-polar residues in the predicted pocket of SanA and SanA function was assayed by EOP. Mutations to the residues on the surface of the pocket affected SanA function, while mutations to residues buried inside SanA did not. **(F)** Mutations were made to residues predicted to be on or near the surface of SanA and SanA function was assayed by EOP. Several mutations exhibited partial loss of function. R69E serves as a control as a pocket mutation with apparent full loss of function. Images are representative of at least three independent experiments.

## DISCUSSION

Here, we set out to understand how SanA affects the cell’s envelope permeability barrier. We confirmed that a Δ*sanA* strain has envelope permeability phenotypes that occur under carbon limitation and high temperatures. Additionally, we discovered that a Δ*sanA* Δ*wecA* strain, which has heightened isoprenoid carrier levels due to a lack of ECA synthesis (40), had a pronounced, synthetic SDS-EDTA sensitivity and that this sensitivity could be suppressed by partial loss of function mutations to either *ftsI* or to *prc*. When investigating the suppression phenotype, we found that mutations to septal PG synthesis machinery, but not early divisome assembly, could suppress the Δ*sanA* Δ*wecA* SDS-EDTA sensitivity. Loss of SanA resulted in decreased PG incorporation in the region of the septum and this was suppressed in dividing cells by the *ftsI* suppressor mutation. Moreover, we observed that overexpressing *murA* in a Δ*sanA* strain phenocopied Δ*sanA* Δ*wecA*, indicating increased lipid II is important for the SDS-EDTA phenotype. Finally, at least one of the *ftsI* suppressor mutations we isolated, *ftsI_V98E_*, resulted in increased FtsW activity, which likely results in the increased septal PG synthesis.

Based on these findings, we propose a model which provides a parsimonious explanation for our results (**Figure 9**). During normal growing conditions, precursors are carefully allocated to allow for balanced synthesis of lipid II and other envelope constituents (**Figure 9A**). In our model, we reason that, under certain cellular stress conditions, the amount of lipid II available for PG synthesis increases. For example, enzymes involved in lipid II synthesis might have higher activity rates at higher temperature leading to increased lipid II synthesis. Alternatively, under carbon-limited conditions, decreased growth and the lower surface area of stationary phase cells could lead to higher lipid II levels. The increase in substrate availability creates a potential imbalance of septal and elongation PG synthesis, which is prevented by SanA (**Figure 9B**). When SanA is absent, lipid II use for lateral PG elongation and septal PG synthesis is imbalanced and this imbalance ultimately leads to envelope permeability (**Figure 9C**). This model would also suggest loss-of-function mutations in early divisome assembly increase the membrane permeability of a Δ*sanA* strain by decreasing the number of active septal PG synthesis complexes and so further decreasing septal PG synthesis rates. In SanA’s absence, mutations that restore septal PG synthesis rates restore balance in septal PG synthesis and so the envelope permeability barrier.

**Figure 9:**
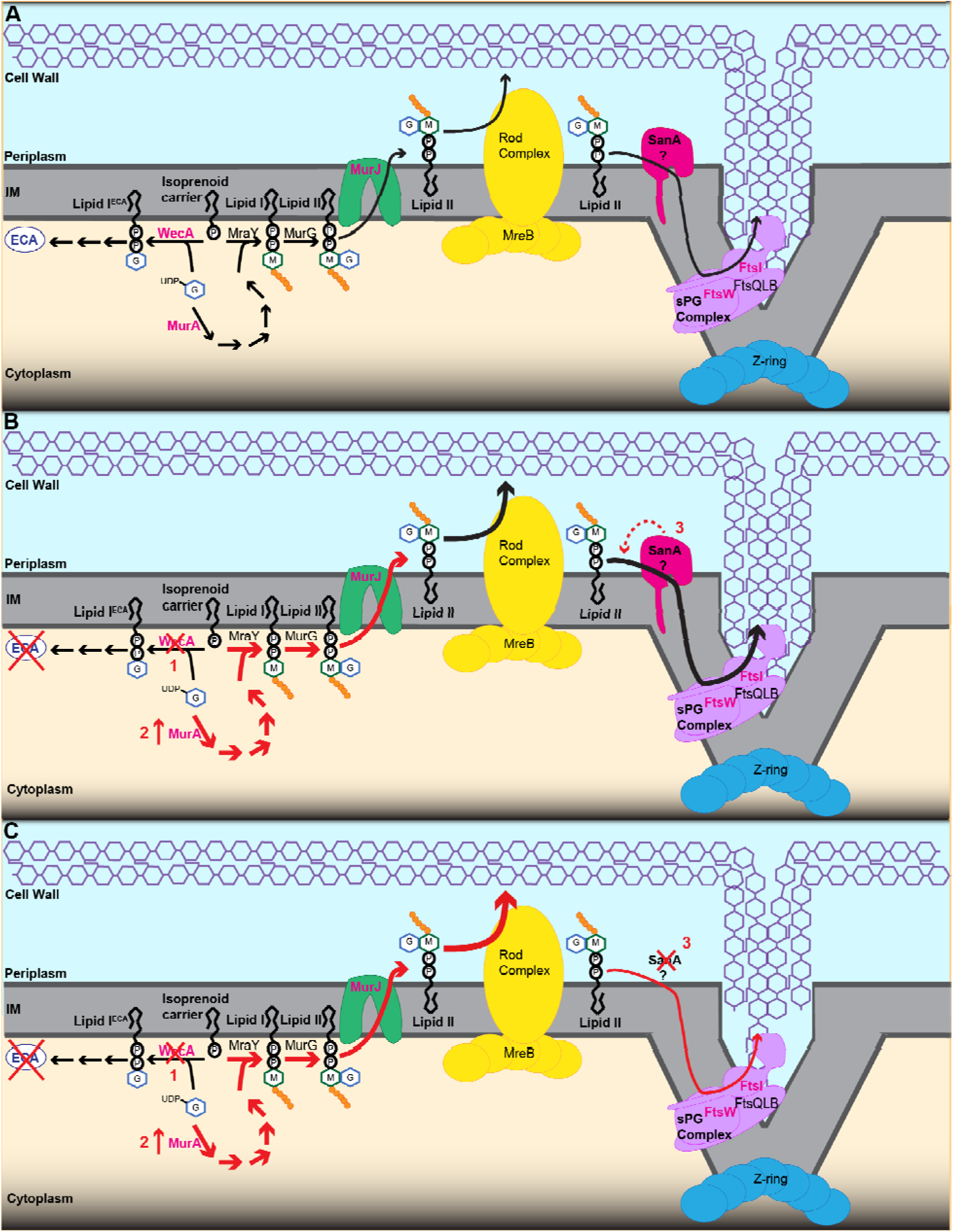
Hypothesized model for the role of SanA in modulating lipid II availability for septal PG synthesis. **(A)** The synthesis of lipid II begins with MurA which catalyzes the first committed step of PG synthesis. The isoprenoid carrier (undecaprenyl-phosphate) is used for the synthesis of extracytoplasmic glycans including ECA and PG. This carrier is allocated between these synthesis pathways to maintain balanced synthesis. Once lipid II is synthesized, it is flipped across the IM by MurJ. Then, the available lipid II is used by the Rod complex for elongating the cell wall and by the septal PG complex for building the septum during cell division. We hypothesize that SanA controls the availability of lipid II to FtsW and FtsI in the septal PG complex. **(B)** Removing WecA prevents use of the isoprenoid carrier for ECA synthesis increasing its availability for the synthesis of other glycans including peptidoglycan (1). Alternatively, overexpressing *murA* increases flux through the lipid II synthesis pathway, increasing lipid II levels. We hypothesize that SanA balances the increased lipid II feeding into septal and lateral PG synthesis, maintaining balanced synthesis of the envelope components. When SanA is absent (3) and lipid II levels are increased, the increased lipid II levels cause increased lateral PG synthesis compared to septal PG synthesis, and this causes increased envelope permeability.

In support of this model, a paper (81) was published while this manuscript was in preparation that identified a suppressive phenotype of Δ*sanA* for the growth of several cell division mutants including *ftsI23* and found that PG incorporation was increased in a Δ*sanA* strain (81). Although the paper examined different conditions from our study, it also found synthetic interactions between early genes in ECA synthesis and Δ*sanA*, specifically for growth in low osmolarity (81). Furthermore, the paper demonstrated that Δ*sanA* phenotypes could be suppressed by endopeptidase overexpression, which provides a possible explanation for our *prc* suppressors, which could increase endopeptidase levels as well as prevent the cleavage of FtsI (24, 81). From these findings it was unclear whether SanA’s phenotypes related to septal PG synthesis or PG synthesis more broadly. However, our work clearly demonstrates that the membrane permeability phenotype, which occurs when increased lipid II levels are coupled with the absence of SanA, specifically relates to septal PG synthesis and that changes to the elongasome (involved in cell lengthening) are not necessary to prevent permeability.

How SanA maintains septal PG synthesis with different lipid II levels remains an unanswered question. The recent paper speculates that SanA may have enzymatic activity, based on phenotypes of several *sanA* point mutations (81). We agree that predicted structures of SanA suggest it has a potential binding pocket and possibly enzymatic activity. Here, we present a large panel of *sanA* point mutations that demonstrate conserved residues within this pocket are important for SanA function. If SanA does have enzymatic activity, it is possible it alters lipid II in some way. Alternatively, SanA may bind to and sequester lipid II from PG elongation. However, SanA molecules in the cell are 10 to 50-fold fewer than lipid II under normal growth conditions (107, 108). Thus, a significant increase in SanA levels under activating conditions would be necessary for this to occur. Lipid II synthesis is regulated through dimerization of MraY which binds to excess lipid II, inhibiting its activity and lipid II synthesis (109). Given this, it is possible SanA is involved in this regulatory pathway. However, it’s perhaps more likely that SanA directly controls the availability of lipid II for septal PG synthesis rather than overall lipid II levels (see **Figure 9**). For instance, SanA could feed lipid II to the septal PG synthesis complex and ensure correct substrate availability to balance septal and lateral PG synthesis. This hypothesis could explain the slight increase in cell length observed in the Δ*sanA* strain, where cells elongate further before division. Finally, a paper was published during revision of this manuscript that suggests direct or indirect physical interaction between SanA and PBP1b (110). Thus, SanA could act at least in part by modulating PBP1b activity. However, differentiating these possibilities will require future investigation.

FtsI’s cytoplasmic region and transmembrane helix are important for its recruitment to the Z-ring through interaction with FtsW (80). The periplasmic portion of FtsI contains the pedestal domain, which is important for interacting with other members of the septal PG synthesis complex, and the catalytic domain that acts as the transpeptidase during septal PG synthesis (80). Both FtsI variants that we identified, FtsI^V98E^ and FtsI^S165P^, are in the pedestal domain of FtsI. Notably, this domain is proposed to recruit FtsN (the final activating protein for the complex), to interact with FtsW (the transglycosylase) and to control its activation, and to control FtsI transpeptidase activity (80, 84, 111). It is interesting then that although both FtsI^V98E^ and FtsI^S165P^ resulted in longer cells, suggesting a partial loss-of-function, FtsI^V98E^ also increased PG incorporation in the area of the septum and made wild type cells resistant to FtsW depletion, suggesting increased septal PG synthesis. Thus, these mutations may affect both FtsI and FtsW activity to alter septal PG synthesis. Mutations to residues near S165 (R166 and R167) have been shown to either cause constitutive activation of FtsW or loss-of-function in FtsI transpeptidase activity (80, 112, 113) so it may be that S165P causes functional changes to both FtsI and FtsW as well. Further investigation of these mutants may lead to greater insight into the complex regulatory interactions carried out by the pedestal domain of FtsI.

Finally, it is worth considering how an imbalance in septal versus lateral PG synthesis could lead to increased envelope permeability. The synthesis of PG and OM components share many precursors. Therefore, it is possible that an increase in elongation PG synthesis could limit some OM components, hindering proper OM synthesis. However, increasing septal PG synthesis should not clear this defect. More likely, the envelope biosynthesis phenotype is due to a lack of coordination between different envelope biosynthetic pathways. For instance, OMP biosynthesis has been shown to increase in the presence of immature PG (48). Therefore, it is possible that decreased septal PG synthesis leads to an decrease in immature PG, causing reduced OMP insertion near the septum. It is also possible that there is a defect in OM invagination caused by the alteration in septal PG synthesis. These questions will be interesting avenues for future investigations.

## MATERIALS AND METHODS

### Strains and growth conditions

All strains used in this study are listed in **Table S1**. Cultures were inoculated, grown at 37°C unless otherwise noted in LB, M63 or MOPS media, and supplemented with 20mg/liter chloramphenicol, 25mg/liter kanamycin, 10mg/liter tetracycline, or a combination thereof as necessary. M63 was supplemented with 1mM MgSO_4_, 50mg/ml thiamine, and 0.2% glucose or 0.05% glucose for carbon-limitation. Mutations were moved using P1*vir* transduction (114) and resistance cassettes were removed with the Flp recombinase-FRT system (115). To clone a plasmid encoding *sanA*, genomic DNA was amplified using the primers pZS21-RBS-*sanA* F and *sanA*-pZS21 R (**Table S2**). pZS21 (116) was amplified using pZS21 F and pZS21-RBS R (**Table S2**). The fragments were assembled using HiFi Assembly Master Mix (New England Biolabs [NEB]) as per the manufacturer’s instructions. Site-directed mutagenesis was performed by PCR with Q5 polymerase (NEB) and the indicated primers (**Table S2**). PCR products were digested with DpnI (NEB), purified, phosphorylated with T4 polynucleotide kinase (NEB), and then self-ligated and transformed. Mutations were confirmed by sequencing.

### Sensitivity assays

Resistance to SDS in stationary phase was assay as previously described (49). For EOPs, overnight cultures were serially diluted, plated on the indicated plates, and plates were incubated overnight. For Kirby-Bauer tests, 100 µL of overnight cultures were added to 3 mL liquified top LB agar and spread on LB agar plates. Preconditioned antibiotic disks were placed on the agar surface and plates were incubated overnight. The diameter of the zone of clearance was measured in at least two directions. For MICs, overnight cultures were normalized to an OD_600_ of 0.1 then further diluted 1:1000. Diluted cultures were then added to 96-well plate followed by two-fold dilutions of the indicated antibiotic. Plates were incubated at 37 °C overnight. Then, OD_600_ was read using a BioTek Synergy H1 plate reader. The lowest concentration where growth is inhibited was determined to be the MIC.

### Lipid A modification assay

Cultures were grown overnight in the indicated media. Cells were then diluted 1:100 in fresh medium containing 2.5 µCi/mL of ^32^P and grown to OD_600_ 0.6-0.8. LPS was purified and mid-acid hydrolysis was used to liberate lipid A which was visualized using thin-layer chromatography (TLC) as previously described (117).

### Suppressor isolation

To isolate suppressors, cultures were serial diluted and plated with 0.5% SDS with 1.0 or 1.1 mM EDTA. Colonies were isolated and suppression was confirmed by EOP. Verified suppressors were categorized based on antibiotic resistance profiles. For a representative of each category and the parent strain, genomic DNA was purified using the DNeasy Blood and Tissue Kit (Qiagen) and provided to SeqCoast Genomics for library preparation (using a DNA Prep Tagmentation Kit (Illumina) and unique dual indexes) and sequencing with an Illumina NextSeq2000 using a 300-cycle flow cell producing 2×150bp reads. A PhiX control was used to optimize base calling. Demultiplexing, read trimming, and run analytics were performed using DRAGEN software version. SeqCoast quality trimmed the reads, mapped them to the MG1655 reference genome (Genebank ID U00096.3) and called variants using Trimmomatic and Breseq software (118–120). Mutations were confirmed in all suppressors using Sanger sequencing. *ftsI* mutations were linked to *zab*-3051:Tn*10* using P1*vir* transduction (121).

### P*_rprA_* reporter activity assay

Overnight cultures carrying the pJW15-P*_rprA_* reporter plasmid (76, 77) were assayed for luciferase activity as has been previously described (70).

### Cell imaging and analysis

Overnight cultures were diluted 1:10,000 and grown to OD_600_ ∼0.5 before imaging. Imaging and cell shape analysis were performed as previously described (122, 123). Differential interference contrast (DIC) images of cells on an agarose pad were captured using an Andor iXon Ultra 897 EMCCD camera (effective pixel size 160 nm) on an inverted Nikon Eclipse-Ti™ microscope with a 100×1.49 NA TIRF objective. Cell length and width was determined using a MATLAB (MathWorks, Inc) script (122). Raincloud plots were made and statistics calculated using Raincloud-shiny (124). For protein localization, overnight cultures were diluted to OD_600_ 0.1 and grown to OD_600_ 0.4. GFP-tagged proteins were excited by a 488 nm laser. GFP-tagged *ftsZ* was expressed from pXY027 (94) (Addgene #98915). Demographs were constructed using ImageJ plug-in MicrobeJ (125). Cells were aligned at mid-cell and sorted by their length, and fluorescence intensity was normalized.

For assaying PG synthesis, cells were grown to exponential phase and then incubated in LB supplemented with 100 µM BODIPY-FL-3-amino-D-alanine (BADA) in LB for 10 minutes at 37 °C. The cells were collected by centrifuging 1 minute at 9,000xg and washed three times with LB before imaging as above. Images were analyzed using code available on Zenodo (https://doi.org/10.5281/zenodo.18839357). Briefly, fluorescence intensity across cells was normalized to their surface area assuming a cylindrical shape with a width of 1 µm. Normalized single cell fluorescence intensity profiles were integrated along the cell axis using trapezoidal numerical integration. Center cell intensity was calculated as the intensity within ±5% of the midpoint of the cell. Cells were classified as dividing or non-dividing based on the z-score of fluorescence enrichment at mid cell (±10% of cell length) with a threshold of z>1.0.

### Immunoblot Analysis

Immunoblots were performed as previously described (126, 127). The primary antibodies used were M2 αFLAG (Millipore Sigma, 1:50,000 dilution) and αGroEL (Millipore Sigma, 1:60,000). α-mouse and α-rabbit secondary antibodies (Prometheus) were used at a 1:100,000 dilution.

## ACKNOWLEDGEMENTS

Thank you to the current and former members of the Mitchell Lab as well as Drs. Jolene Ramsey, Jennifer Herman, Deborah Siegele, Ryland Young (Texas A&M University), Anna Konovalova (University of Texas Health Science Center Houston), Petra Levin (Washington University in St. Louis), and Natividad Ruiz (The Ohio State University) for insights and productive discussions. Thank you to Dr. Srutha Venkatesan for helping with the initial stages of the cell-length experiments. We would also like to thank Dr. Thomas Silhavy (Princeton University) for all his assistance with the early stages of this project. Thank you to Drs. Petra Levin (Washington University in St. Louis), Natividad Ruiz (The Ohio State University), and Thomas Bernhardt (Harvard University Medical School) for sharing strains with us to facilitate these experiments and to Dr. Tracy Raivio (University of Alberta) for providing the pJW15-P*_rpra_* reporter plasmid and Dr. Jie Xiao (Johns Hopkins University School of Medicine) for providing pXY027 (Addgene #98915).

This work was supported by the National Institute of Allergy and Infectious Diseases under award numbers R01-AI155915 (to A.M.M.) and R01-AI176766 (to M.S.T.), and the National Institute of General Medical Sciences under award number R01-GM129000 (to B.N.).

