## Supplemental Material for "Impaired envelope integrity in the absence of SanA is linked to increased lipid II availability and an imbalance of septal peptidoglycan synthesis"

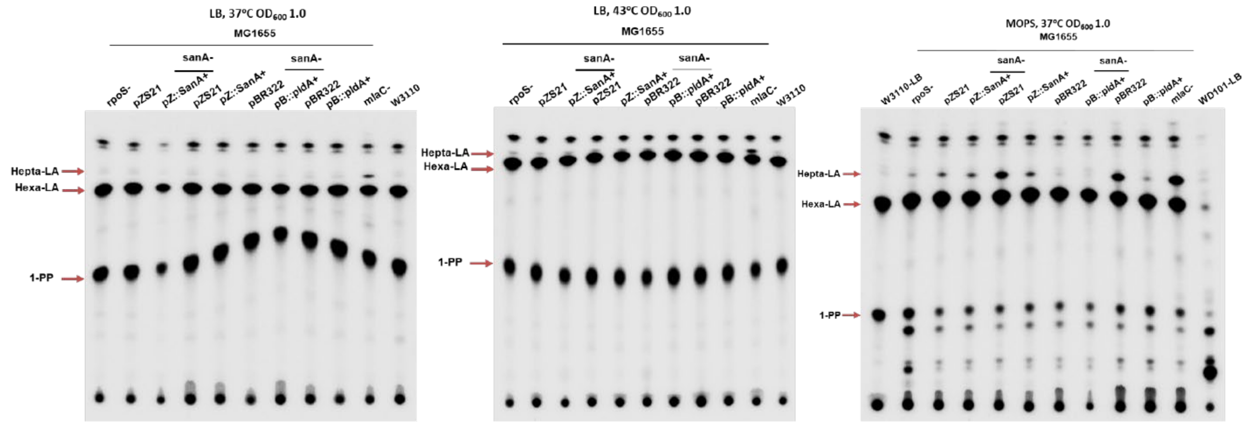


**Figure S1: Loss of SanA causes accumulation of hepta-acylated lipid A when cells are grown with glucose as a carbon source.** Full TLC images are shown for the experiment shown in Figure 1D.

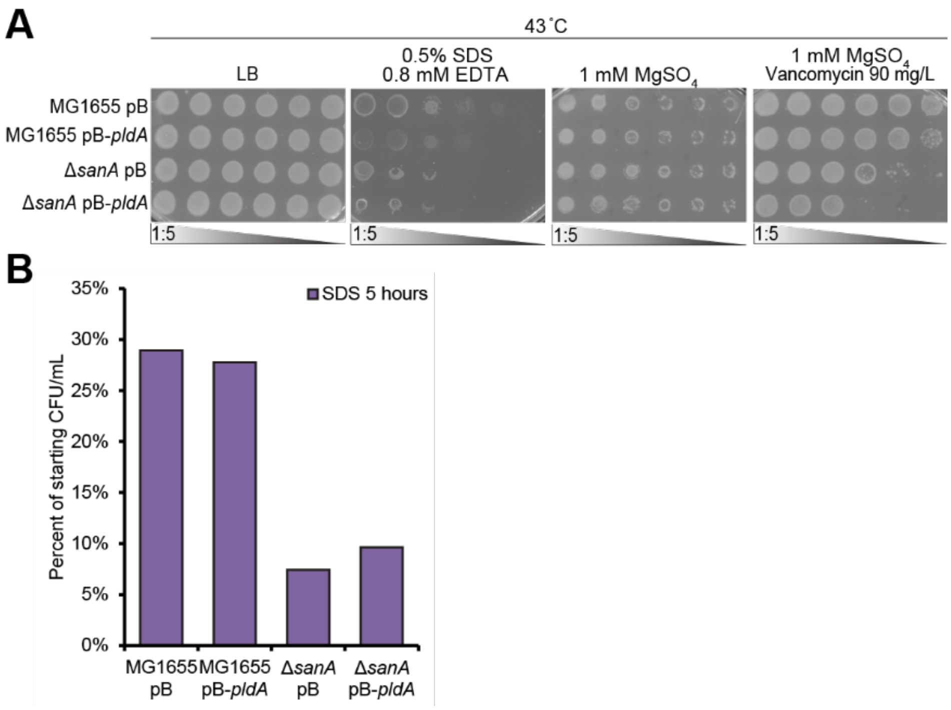


**Figure S2: Overexpression of *pldA* does not suppress the envelope permeability phenotypes caused by loss of SanA. (A)** The effect of *pldA* overexpression on the SDS-EDTA resistance and vancomycin resistance of a Δ*sanA* strain at 43 °C was assayed by EOP. Overexpression of *pldA* did not suppress these phenotypes. **(B)** The viability of the indicated strains was assayed after 5 hours of SDS treatment during carbon-limitation stationary phase. Overexpression of *pldA* did not restore the SDS resistance of the Δ*sanA* strain.

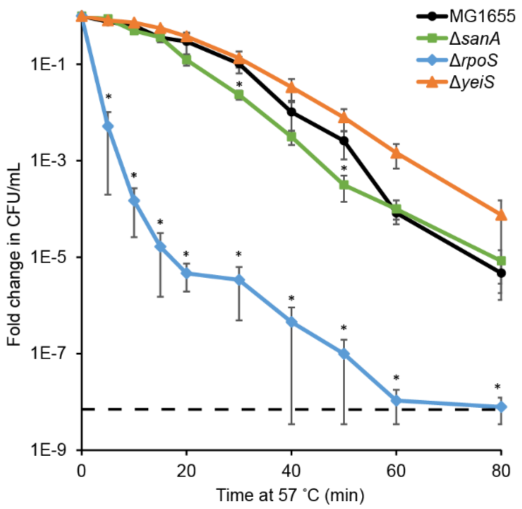


**Figure S3: Temperature sensitivity caused by loss of RpoS is greater than that caused by loss of SanA.** The Δ*sanA* strain grown to carbon limitation has decreased heat shock survival at 57 °C compared to wild type, but less so than an RpoS mutant. This is not true of *sanA* operon partner *yeiS* which does not show temperature sensitivity. Data for the wild type and Δ*sanA* strains are the same as is shown in Figure 1E. Data are the average of 3 to 4 biological replicates ± the SEM. * $p<0.05$ by the Mann Whitney Test.

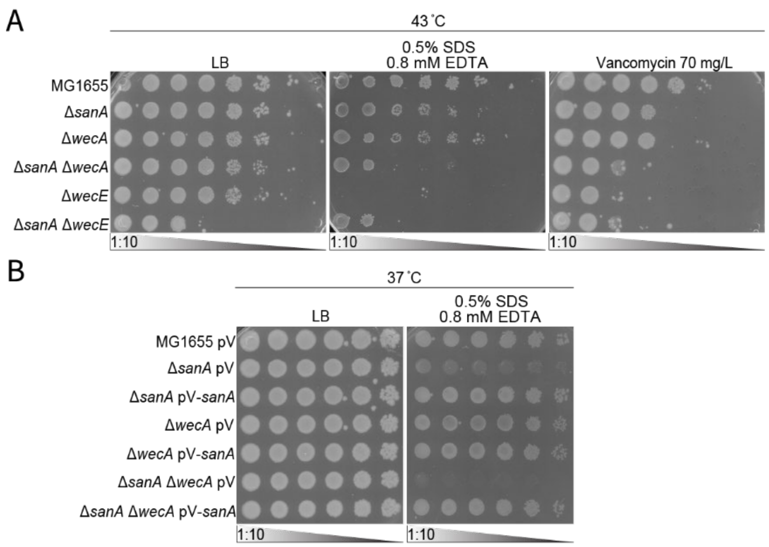


**Figure S4: *sanA* has genetic interactions with mutations in ECA synthesis. (A)** The SDS-EDTA sensitivity of a Δ*sanA* Δ*wecA* strain is maintained at 43 °C; however, a vancomycin phenotype is less apparent. **(B)** The SDS-EDTA sensitivity of a Δ*sanA* Δ*wecA* strain can be complemented by plasmid-based expression of *sanA*. **(C)** $OD_{600}$ growth curve corresponding to the reporter assay in Figure 2C. Data are shown as the average of three biological replicates ± the SEM.

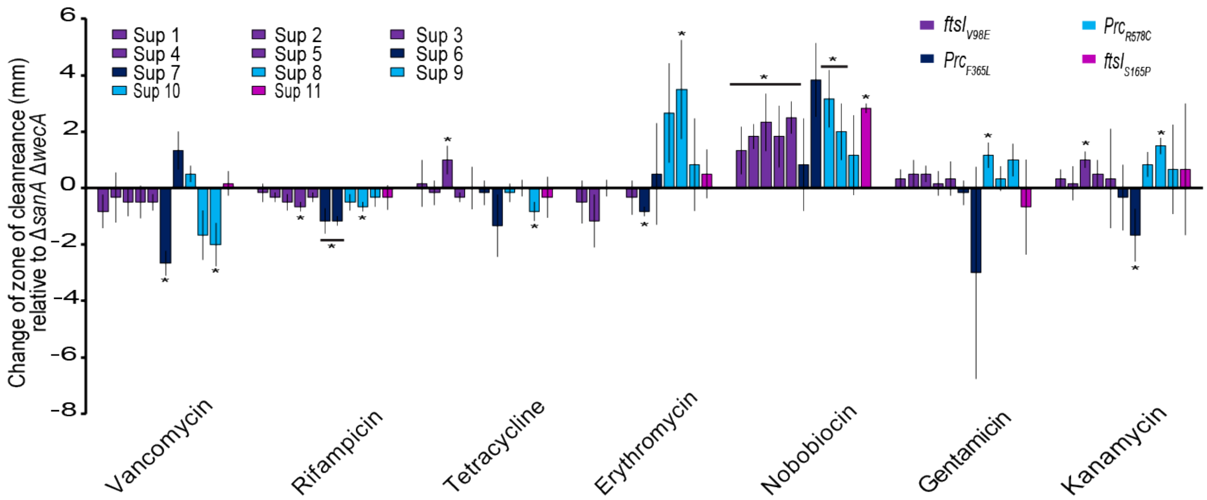


**Figure S5: Suppressors have differing antibiotic resistance profiles.** The isolated suppressor mutations were assayed by Kirby-Bauer tests for their resistance to a panel of antibiotics. Data are shown as the change in zone of clearance relative to the Δ*sanA* Δ*wecA* strain. For ease of viewing, suppressors have been renumbered and grouped by phenotype. The color of the bars indicates the mutation identified in the indicated strain. Data are the average of three biological replicates ± the SEM.

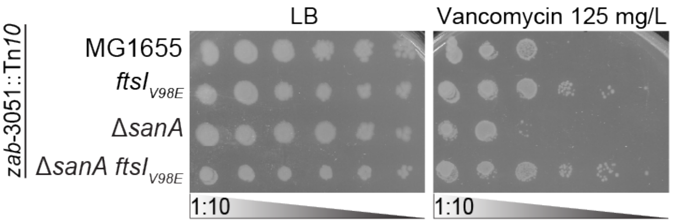


**Figure S6: FtsI$^{V98E}$ causes increased vancomycin resistance.** EOPs were performed at 43 °C to determine the vancomycin resistance of the indicated strains. The *ftsI$_{V98E}$* mutation caused increased vancomycin resistance in both the wild-type and Δ*sanA* backgrounds. Images are representative of three independent experiments.

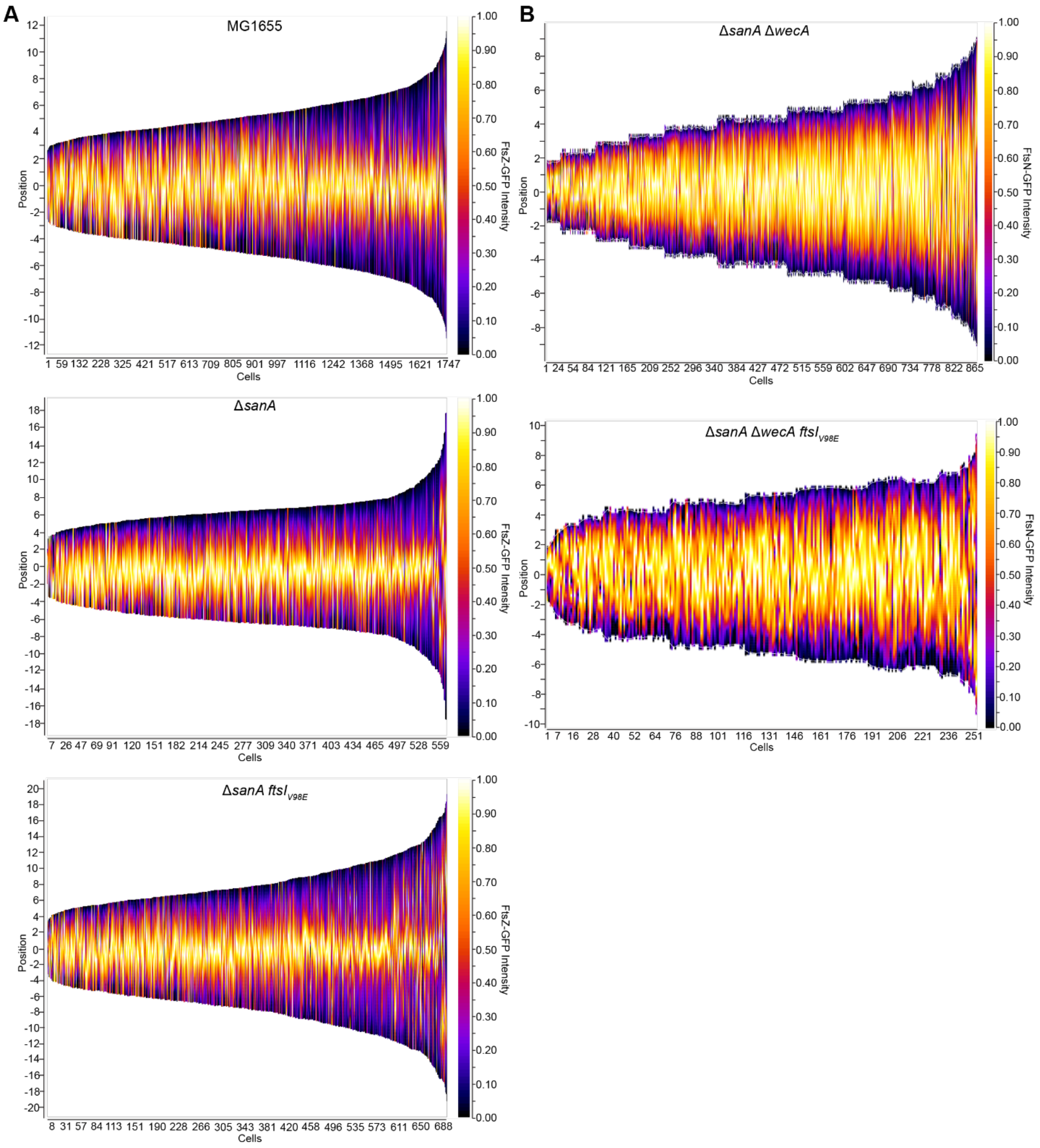


**Figure S7: Localization of FtsZ and FtsN do not correlate with phenotypes associated with loss of SanA. (A)** The FtsZ-GFP localization was assayed in the wild-type strain, the Δ*sanA* strain, and the Δ*sanA* strain carrying the $ftsI_{V98E}$ mutation. Demographs were computed using the locational data with cells arranged from shortest to longest. No significant differences in FtsZ localization were observed. **(B)** FtsN-GFP localization was assessed in the Δ*sanA* Δ*wecA* and Δ*sanA* Δ*wecA* $ftsI_{V98E}$

strains. Demographics were computed using the localization data. No significant differences in FtsN localization were observed.

**Table S1: Strains used in this study**

| Strain | Genotype | Reference | Notes |
|---|---|---|---|
| MG1655 | K-12 $F^-$ $\lambda^-$ *rph-1* | (1) | CGSC 6300 |
| AM104 | MG1655 Δ*rpoS* | (2) | |
| AM158 | MG1655 Δ*sanA* | (2) | |
| AM202 | MG1655 pZS21 | This study | Plasmid from (3) |
| AM203 | MG1655 pZS21-*sanA* | This study | |
| AM213 | MG1655 Δ*sanA* pZS21 | This study | Plasmid from (3) |
| AM214 | MG1655 Δ*sanA* pZS21-*sanA* | This study | |
| AM237 | MG1655 Δ*yeiS* | This study | |
| AM334 | MG1655 Δ*wecA* | (4) | |
| AM337 | MG1655 Δ*wecE* | (4) | |
| AM363 | MG1655 Δ*mlaC* | This study | |
| AM380 | MG1655 pBR322 | This study | Plasmid from (5) |
| AM381 | MG1655 pBR322-*pldA* | This study | Plasmid from (5) |
| AM382 | MG1655 Δ*sanA* pBR322 | This study | Plasmid from (5) |
| AM383 | MG1655 Δ*sanA* PBR322-*pldA* | This study | Plasmid from (5) |
| AM384 | MG1655 pBR322-*pldA* | This study | Plasmid from (5) |
| AM771 | MG1655 Δ*sanA* Δ*wecA* | This study | |
| AM772 | MG1655 Δ*sanA* Δ*wecE* | This study | |
| AM783 | MG1655 Δ*wecF* | This study | |
| AM784 | MG1655 Δ*wecG* | This study | |
| AM785 | MG1655 Δ*sanA* Δ*wecF* | This study | |
| AM786 | MG1655 Δ*sanA* Δ*wecG* | This study | |
| RB006 | MG1655 Δ*sanA* Δ*wecA* $ftsI_{V98E}$ | This study | Genome nt 91705 T→A |
| RB007 | MG1655 Δ*sanA* Δ*wecA* $ftsI_{S165P}$ | This study | Genome nt 91905 T→C |
| RB010 | MG1655 Δ*sanA* Δ*wecA* $prc_{F365L}$ | This study | Genome nt 1913722 G→C |
| RB011 | MG1655 Δ*sanA* Δ*wecA* $prc_{R578C}$ | This study | Genome nt 1913085 G→A |
| RB036 | MG1655 *zab-3051*::Tn*10* | This study | |
| RB037 | MG1655 $ftsI_{V98E}$ *zab-3051*::Tn*10* | This study | |
| RB038 | MG1655 Δ*sanA* $ftsI_{V98E}$ *zab-3051*::Tn*10* | This study | |
| RB039 | MG1655 Δ*wecA* *zab-3051*::Tn*10* | This study | |
| RB040 | MG1655 Δ*wecA* $ftsI_{V98E}$ *zab-3051*::Tn*10* | This study | |
| RB041 | MG1655 Δ*sanA* *wecA* *zab-3051*::Tn*10* | This study | |
| RB044 | MG1655 Δ*sanA* *wecA* $ftsI_{V98E}$ *zab-3051*::Tn*10* | This study | |
| DW012 | MG1655 Δ*sanA* *wecA* pCH201 (*ftsN*-GFP) | This study | Plasmid from (6) |
| DW014 | MG1655 Δ*sanA* *wecA* *ftsI23* *leu82*::Tn*10* | This study | *ftsI* allele from (7) |
| DW017 | MG1655 Δ*sanA* *wecA* *ftsA** *leu82*::Tn*10* | This study | *ftsA* allele from (8) |
| DW041 | MG1655 pXY027 | This study | Plasmid from (9) |
| DW042 | MG1655 Δ*sanA* pXY027 | This study | Plasmid from (9) |
| DW043 | MG1655 Δ*sanA* Δ*wecA* pXY027 | This study | Plasmid from (9) |

| Strain | Genotype | Reference | Notes |
|---|---|---|---|
| DW068 | MG1655 Δ*sanA wecA ftsZ*84 *leu*::Tn*10* | This study | *ftsZ* allele from (10) |
| DW069 | MG1655 Δ*sanA ftsZ*84 *leu*::Tn*10* | This study | *ftsZ* allele from (10) |
| DW070 | MG1655 Δ*sanA ftsA12 leu82*::Tn*10* | This study | *ftsA* allele from (11) |
| DW071 | MG1655 Δ*sanA ftsA27 leu-260*::Tn*10* | This study | *ftsA* allele from (12) |
| DW072 | MG1655 Δ*sanA wecA ftsL* leu82*::Tn*10* | This study | *ftsL* allele from (13) |
| DW073 | MG1655 Δ*sanA wecA ftsQ1 leu82*::Tn*10* | This study | *ftsQ* allele from (14) |
| DW074 | MG1655 Δ*sanA wecA ftsA27 leu-260*::Tn*10* | This study | *ftsA* allele from (12) |
| DW075 | MG1655 Δ*sanA wecA ftsA12 leu82*::Tn*10* | This study | *ftsA* allele from (11) |
| DW076 | MG1655 Δ*sanA ftsL* leu82*::Tn*10* | This study | *ftsL* allele from (13) |
| DW077 | MG1655 Δ*sanA ftsQ1 leu82*::Tn*10* | This study | *ftsQ* allele from (14) |
| DW081 | MG1655 *ftsZ*84 *leu*::Tn*10* | This study | *ftsZ* allele from (10) |
| DW083 | MG1655 *ftsA12 leu82*::Tn*10* | This study | *ftsA* allele from (11) |
| DW084 | MG1655 *ftsA27 leu-260*::Tn*10* | This study | *ftsA* allele from (12) |
| DW085 | MG1655 Δ*ftsW* Δ*araH zab*-3051::Tn*10* pCS37 | This study | Plasmid and *ftsW* allele from (15) |
| DW089 | MG1655 Δ*ftsW* Δ*araH zab*-3051::Tn*10* $ftsI_{V98E}$ pCS37 | This study | Plasmid and *ftsW* allele from (15) |
| DW091 | MG1655 Δ*sanA* Δ*ftsW* Δ*araH zab*-3051::Tn*10* $ftsI_{V98E}$ pCS37 | This study | Plasmid and *ftsW* allele from (15) |
| DW092 | MG1655 Δ*sanA* Δ*wecA* Δ*ftsW* Δ*araH zab*-3051::Tn*10* $ftsI_{V98E}$ pCS37 | This study | Plasmid and *ftsW* allele from (15) |
| DW095 | MG1655 Δ*sanA* Δ*wecA* Δ*ftsW* Δ*araH zab*-3051::Tn*10* pCS37 | This study | Plasmid and *ftsW* allele from (15) |
| DW096 | MG1655 *ftsL* leu82*::Tn*10* | This study | *ftsL* allele from (13) |
| DW097 | MG1655 *ftsQ1 leu82*::Tn*10* | This study | *ftsQ* allele from (14) |
| DW099 | MG1655 Δ*sanA* Δ*wecA murJ* Ω(−14::*bla araC* $P_{BAD}$) Δ*araH zab*-3051::Tn*10* | This study | *murJ* allele from (16) |
| DW0101 | MG1655 *murJ* Ω(−14::*bla araC* $P_{BAD}$) Δ*araH zab*-3051::Tn*10* Δ*sanA* | This study | *murJ* allele from (16) |
| DW102 | MG1655 *murJ* Ω(−14::*bla araC* $P_{BAD}$) Δ*araH zab*-3051::Tn*10* | This study | *murJ* allele from (16) |

| Strain | Genotype | Reference | Notes |
|---|---|---|---|
| DW103 | MG1655 *murJ* Ω(−14::*bla araC* $P_{BAD}$) Δ*araH zab*-3051::Tn*10* Δ*sanA* Δ*wecA ftsI*$^{V98E}$ | This study | *murJ* allele from (16) |
| DW104 | MG1655 *murJ* Ω(−14::*bla araC* $P_{BAD}$) Δ*araH zab*-3051::Tn*10* Δ*sanA ftsI*$^{V98E}$ | This study | *murJ* allele from (16) |
| DW108 | MG1655 Δ*sanA* Δ*wecA zab*-3051::Tn*10 ftsI*$^{V98E}$ pCA24N-*murA* | This study | Plasmid from (17) |
| DW109 | MG1655 Δ*sanA zab*-3051::Tn*10 ftsI*$^{V98E}$ pCA24N-*murA* | This study | Plasmid from (17) |
| DW110 | MG1655 Δ*sanA* Δ*wecA zab*-3051::Tn*10* pCA24N-*murA* | This study | Plasmid from (17) |
| DW111 | MG1655 Δ*sanA zab*-3051::Tn*10* pCA24N-*murA* | This study | Plasmid from (17) |
| DW112 | MG1655 *zab*-3051::Tn*10* pCA24N-*murA* | This study | Plasmid from (17) |
| DW113 | MG1655 *zab*-3051::Tn*10* pCA24N | This study | Plasmid from (17) |
| DW114 | MG1655 Δ*sanA* Δ*wecA zab*-3051::Tn*10* pCA24N | This study | Plasmid from (17) |
| DW115 | MG1655 Δ*sanA zab*-3051::Tn*10* pCA24N | This study | Plasmid from (17) |

**Table S2: Primers used in this study**

| Primer | Sequence (5' to 3') |
|---|---|
| pZS21-RBS-*sanA* F | gtatcgataagcttgatatcaggaggacagctATGTTAAAGCGCGTGTTC |
| *sanA*-pZS21 R | gatcccccgggctgcaggaattcCTTTCCTTGTTTCTTTTGTAATTC |
| pZS21 F | GAATTCCTGCAGCCCGGG |
| pZS21-RBS R | AGCTGTCCTCCTGATATCAAGCTTATCGATACCGTCGACCTC |
| sanA-GS-Flag F | tataaagatgatgatgataaatgatgaTCTAGAGGCATCAAATAAAAC |
| sanA-GS-Flag R | atcgccgctacccccctccagagccaccCTTTCCTTGTTTCTTTTGTAATTC |
| sanA_V47-G52_R | TGGCGGTAGGGGAGATCC |
| sanA_V49T_F | GGTCGGTacGGTGCTCGGAACAG |
| sanA_V50T_F | GGTCGGTGTGacGCTCGGAACAG |
| sanA_L51N_F | GGTCGGTGTGGTGaaCGGAACAG |
| sanA_L51A_F | GGTCGGTGTGGTGgcCGGAACAG |
| sanA_G52S_F | GGTCGGTGTGGTGCTCaGtACAG |
| sanA_R69_R | TACTGATTAATTACGCCAG |
| sanA_R69A_F | TTATCGCTACgccATTCAAGGAGC |
| sanA_R69E_F | TTATCGCTACgaaATTCAAGGAGCG |
| sanAE98-M102_R | TAACTTTGCAATGCGTTATCGC |
| sanA_E98A_F | TAATGcGCCGATGACCATGCG |
| sanA_V111-V117_R | CCAGCAGCGATTAAATCTTTG |
| sanA_I116S_F | TGTCGACCCATCAGATtcTGTTC |
| sanA_L118-L126_R | CAATATCTGATGGGTCGACACC |
| sanA_L118N_F | TTaatGATTACGCAGGCTTTCGTACGCTGGA |
| sanA_V130-V135_R | AGCGTACGAAAGCCTGCG |
| sanA_V130T_F | GGATTCCATCacgCGTACACGCAAAGTTTTC |
| sanA_V135T_F | GGATTCCATCGTGCGTACACGCAAAacTTTC |
| sanA_R131-F136_R | ACGATGGAATCCAGCGTAC |
| sanA_F136Q_F | GCGTACACGCAAAGTTcaaGATA |
| sanA_T145_R | AAATCATTAGTATCGAAAACTTTG |
| sanA_T145A_F | CATTATTATCgctCAACGTTTCC |
| sanA_T145N_F | CATTATTATCaatCAACGTTTCCACTGTG |
| sanA_H149-R152_R | CGTTGGGTGATAATAATGAAATCATTAG |
| sanA_H149A_F | TTTCgccTGTGAGCGAGCATT |
| sanA_R152A_F | TTTCCACTGTGAGgcaGCATT |
| sanA_R152E_F | TTTCCACTGTGAGgaaGCATT |
| sanA_L176_R | GGTGACGGTACGGCATAA |
| sanA_L176N_F | GAAAGATATGaatTCAGTACGTATTCGTGAATTTGCC |
| sanA_I180-F183_R | ACTGACAGCATATCTTTC |
| sanA_E182R_F | ACGTATTCGTaggTTTGCCGCCC |
| sanA_F183Q_F | ACGTATTCGTGAAcaaGCCGCCC |
| sanA_D192_R | CCGAAACGGGCGGCAAAT |
| sanA_D192R_F | TGCGCTGGCTcgcCTTTATATTTTTAAACGTG |
| sanA_R198_R | TAAAGGTCAGCCAGCGCA |
| sanA_R198D_F | TATTTTTAAAgacGAACCGCGTTTTTTAGGGCCGC |
| sanA_D192R_R | TAAAGgcgAGCCAGCGCA |
